# Quantifying Effects of Dataset Size, Data Variability, and Data Curvature on Modelling of Simulated Age-Related Motor Development Data

**DOI:** 10.1101/2023.05.07.539755

**Authors:** Stephan C.D. Dobri, Stephen H. Scott, T. Claire Davies

## Abstract

Motor development in children and youth occurs non-linearly; improvements are rapid at younger ages and decrease as they reach adulthood. There is also evidence that performance variability changes as children and youth age. Accurate models of typical performance are necessary to identify deficits in motor performance and to track the efficacy of therapies. Robotic devices have been used previously to measure motor performance in children and youth, and produce models of typical performance; however, power analyses on these models have not been explored.

An algorithm was created to generate normative models of typical motor performance. The accuracy and repeatability of the algorithm were tested using simulated data that changed the number of data points, and the curvature and variability of the data. Two-hundred and eighty-eight participants who are typically developing (ages 5-18) completed a robotic point-to-point reaching task with the Kinarm Exoskeleton. Exponential curves were fit to reaction time measured by the Kinarm to model typical performance. The results of the simulations were used to generate confidence intervals on the models of typical performance.

The simulations showed that number of datapoints had the largest impact on accuracy and repeatability of the models, and that repeatability was age-dependent. The simulations with the uniform and non-uniform datasets generated different confidence intervals; however, these differences were minimal when the number of datapoints at each age were matched between the two datasets.

To ensure identification of deficits is accurately determined, there is a need to account for differences in repeatability when developing models of typical motor performance in children and youth. The results of our simulations can be used to assess repeatability of non-linear models of motor performance based on dataset size in the future.

## Introduction

Children and youth experience motor development in a manner that quickly evolves at younger ages, and the rate of improvement slows until they have reached “near-adult” performance [1-3]. For example, proprioception reaches “near-adult” performance around 8 years of age, whereas other aspects of coordination, such as reaction time, do not mature until 12-14 years of age [1, 4, 5]. Like height and weight, measures of motor performance (e.g. reaction time) of typically-developing children and youth can be plotted against age to generate curves that reflect developmental expectations of motor function. These models of typical development are often used to identify deficits in performance in various patient populations [6-11] based on a comparison to some performance interval of a healthy control population (i.e. 95% of the range of healthy performance or mean +/-2 standard deviations). Therefore, a key first step in this process is the generation of reliable and reproducible curves of developmental behaviour, to accurately identify deficits in performance.

The generation of these developmental curves, including the mean and variance of performance at different ages, can be affected by various factors including the underlying variability of the data and the number of participants included in the analyses. There is evidence that variability in motor performance changes with age [1]. For example, as children reach “near-adult” performance, the variability in their movement decreases [1]. Therefore, using a fixed performance interval width for the whole curve (i.e. one that is calculated based on the entire sample mean) may result in a performance interval that is over-estimated for older ages. This could result in the misidentification of deficits when comparing behavior from patient groups. As well, although it is recognized as good statistical practice, very few studies tend to include power analyses a priori to determine the appropriate number of participants required to achieve desired power [9]. The use of power analyses is particularly pertinent to studies of motor development in light of the non-linearity of age-related performance. For example, studies that recruit participants at older age groups may not require as many participants to achieve sufficient power to identify differences compared to those that recruit younger children. Finally, previous work has highlighted the importance of confidence intervals on fitted curves in order to determine the reliability of the modelled behavioural fits [9]. Further, including confidence intervals at each age along the curves creates confidence bounds which makes it possible to reliably identify deficits in individual patient performance.

The primary objective of the current work was to model non-linear motor performance, accounting for reduced variability with increasing age and to determine the effects of the number of data points included. The secondary objective was to evaluate repeatability using simulated data to create confidence intervals on age-dependent developmental curves. The models within this study provide confidence intervals that change with age to better identify parameters that may indicate deficient performance. Finally, since it is not always possible to collect equal numbers of participants in each age group to create a uniform age distribution, we also wanted to compare uniform vs non-uniform age distributions.

## Materials and Methods

Non-linear modeling of motor development included curve fitting of both uniform and non-uniform distributions, followed by an assessment of accuracy and repeatability. While the uniform data was developed using simulations, the non-uniform dataset represented an age distribution based on data collected from a robotic point-to-point reaching task in which two-hundred and eighty-eight participants who are typically developing (ages 5-18) participated. The non-uniform simulations allowed us to determine if the uniformity of the input data impacted the accuracy and repeatability of the algorithm. Finally, an exponential curve was fit to the performance data collected from the children which included variable confidence intervals based on the simulations.

### Simulated Data with Uniform Distributions

A curve fitting algorithm was developed for non-linear regression models. While simulating changes to the underlying curvature based on the age at which near-adult performance is achieved, effects of sample size and data variability were considered in determining the accuracy and repeatability of the algorithm.

Simulated curves were created with an equal number of datapoints at each integer age, Gaussian white noise was added, and exponential curves were fit to the simulated data. The simulations changed the input curves, dataset sizes, and standard deviation of the Gaussian white noise. Two types of curves were chosen to reflect previously published data on motor coordination of reaching movements: asymptotic growth and asymptotic decay [2, 11, 12]. For example, the path smoothness of a reaching motion in children follows a growth curve with age, whereas reaction time follows a decay curve [2, 11, 12]. The dataset sizes ranged from 14 to 1,792 datapoints to represent small studies and to surpass the sample size of many studies that quantify sensorimotor or cognitive function in children and youth [2, 5, 9, 10, 13, 14]. The standard deviations of the Gaussian white noise (i.e. input SDs) were chosen to reflect reported standard deviations of children and youth performance on motor and proprioception tasks [2, 3, 13, 15-18].

The input curves increased or decreased asymptotically to a value of 1 from 0.5 or 2. The rates of growth or decay reflected different developmental trajectories where data reached “near-adult” performance by age 8, 10, 12, or 14 years. A base curve, represented by the equation below, was used to generate simulated data for participants from ages 5-18:

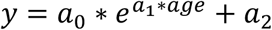

The variable “age” represents the integer age. The coefficients a0, a1, and a2 were calculated to simulate the desired ages of “near-adult” performance. “Near-adult” performance was defined as the curve being within 5% of the final value (0.95 for growth curves, 1.05 for decay curves). Given these parameters, eight input curves, eight dataset sizes, and 15 standard deviations for the Gaussian white noise were used to total 960 simulation combinations. Ten-thousand curve fits were applied for each simulation combination. For each curve fit, new simulated data were created by adding a different random sample of Gaussian white noise to the input curve. For each combination, the 95-percentile range of fitted curves were calculated across the age range to generate confidence intervals for the curve fit.

The MATLAB function “nlinfit.m” was used to fit a non-linear multivariable function to the simulated data (Mathwork, Inc., Massachusetts, USA). The function requires the input data, initial estimations of the coefficients, the independent variable(s) (age in this case), and the desired equation used for curve fitting. After a curve was fit to the data, the standard deviation of the residuals relative to the fitted curve was calculated.

### Simulated Data with Non-Uniform Distributions

Typically, a study does not have an equal distribution of integer ages and sex among the participants. To examine the effects of different numbers of participants, a representative sample of 288 child participants was simulated as non-uniform data. The same process was followed as mentioned above, but using the distribution of participants as shown in Fig. 1. The chosen dataset was based on data that had been collected using a Visually Guided Reaching (VGR) task in the Kinarm robot. The coefficients for creating the input curves, the dataset sizes, and standard deviations can be found in the Supplementary Material.

**Figure 1:**
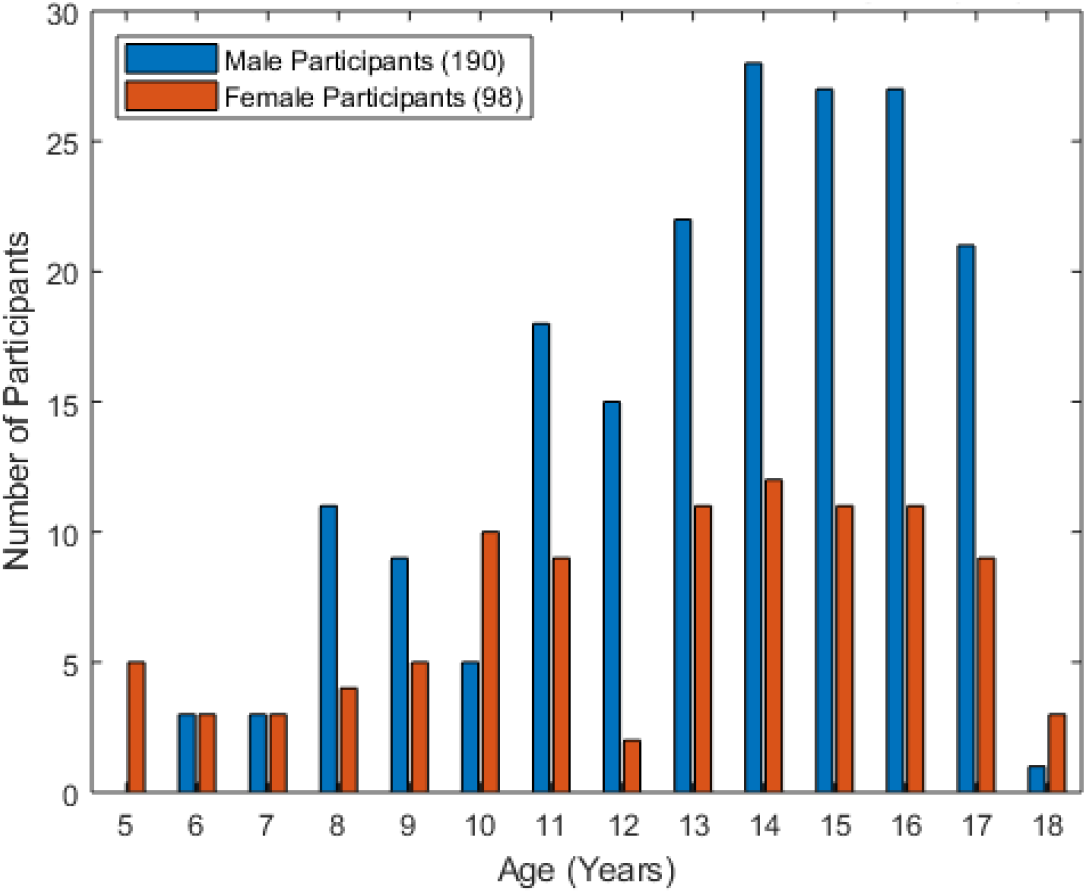
Histogram of age of typically developing children who participated. Nearly twice as many participants were male as were female. Most participants were age 11 or older.

All participants gave informed assent and informed consent was obtained from the guardians prior to participation. This study was approved by the Health Sciences and Affiliated Hospitals Research Ethics Board at Queen’s University, Kingston, Ontario, Canada (application number 6004951) in accordance with the Helsinki Declaration of 1975, as revised in 2000 and 2008.

### Applying Confidence Intervals from both uniform and non-uniform data distributions to data collected from child participants

Data was collected using a Visually Guided Reaching (VGR) task in the Kinarm Exoskeleton Lab (Kinarm, Kingston, Ontario, Canada). Participants sat in a system-integrated modified wheelchair base with their arms supported against gravity. The robot allows free planar movement of the upper limbs while recording elbow and shoulder joint kinematics. Hand position and kinematics were calculated from the joint kinematics. Participants sat in front of a virtual reality display, with vision of their hands and arms occluded, as seen in Fig. 2A. Hand position feedback was displayed in each task on the virtual reality display. The robot has been described in detail previously [19].

**Figure 2:**
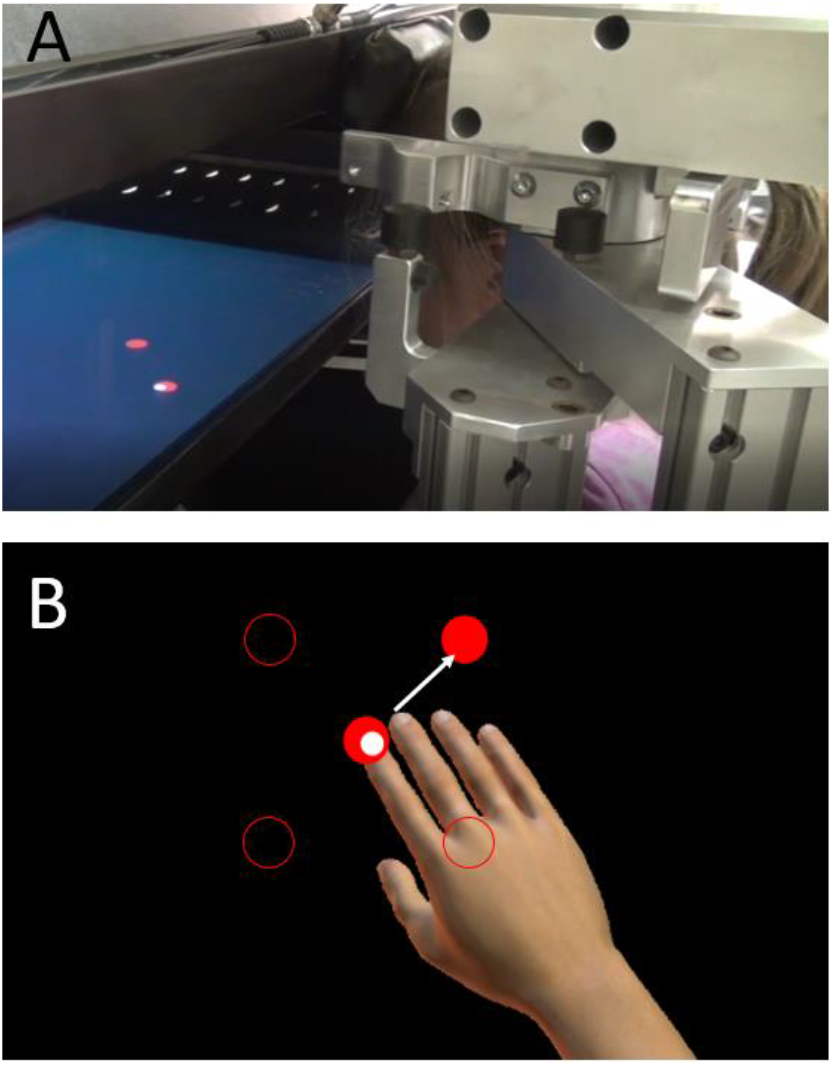
The Kinarm Exoskeleton Robot and representations of Visual Guided Reaching (VGR) task. A) Participant completing a task. Tasks were projected onto a semi-transparent glass sheet from a screen above the glass. Vision of the hands and arms were occluded for all tasks; however, tasks provided visual feedback of hand position. B) VGR task in which the participant was waiting in the central target for a peripheral target to appear. The top-right target appeared so the participant must move the white dot representing hand position into the red target. The positions of the other peripheral targets are represented by the red circle outlines. The image of the participant’s hand was added for clarity.

The Visually Guided Reaching (VGR) task assesses single handed goal-directed reaching [20]. There are four peripheral targets set in a square, 6 cm apart, with a fifth starting target in the centre of the square. Participants are instructed to reach as quickly and accurately as possible to each target as they appear. The peripheral targets appear in a pseudo-random order, six times each for a total of 24 reaches per arm. Each reach consists of moving out to the peripheral target, then back to the central target when it reappears. Participants complete the task once with both their dominant and non-dominant hands. This task is represented in Fig. 2B. In this case, the task is assessed using Reaction Time (RT) [20]. Reaction Time (RT) is defined as the time (in seconds) from when the peripheral target appears to when hand movement onset occurs.

## Results

### Simulated Data with Uniform Distributions

Fig. 3A and 3B display growth and decay input curves when no noise is added to the data. The four different curves in each represent the growth or decay to ages 8, 10, 12, and 14. Fig. 3C-F show how variations in standard deviation (SD) alter the dispersion of data for the growth curve associated with near-adult performance at age 8.

**Figure 3:**
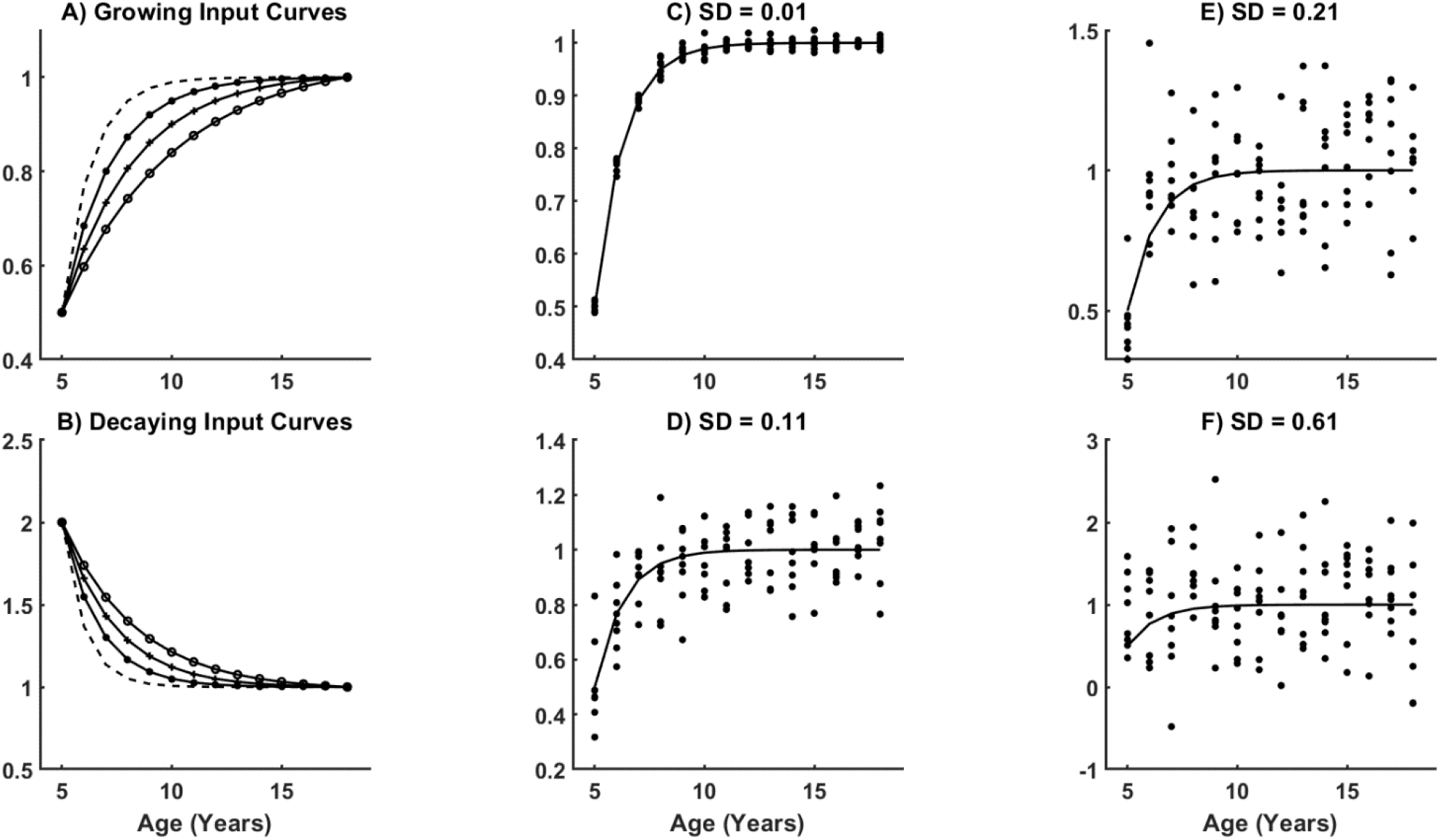
Visualization of all input curves and four different SDs with one input curve. A) shows all growing input curves, whereas B) shows all decaying input curves. C)-F) were all generated with the input curve which grows to “near-adult” performance at age 8, with different input SDs. It is important to note that the scale of each y-axis is fit to the data represented and that they are not the same between any of the above plots. Additionally, the y-axes have been left unlabeled as the data represented are simulated, dimensionless values. The x-axis for all plots is age in years.

Fig. 4A highlights the range of model fits for 10,000 runs for the early development curve (growth to age 8), with a SD of 0.21 and a sample size of 56 datapoints. The estimated mean and associated 95% confidence interval are plotted for each age, as well as the estimated mean +/-2SD and its 95% confidence interval (CI). The estimated 2SD around the mean represents 95% of healthy performance range and can be used as a cut-off value. If an individual datapoint lies outside this 2SD range around the mean, the participant would be identified as having a deficit.

**Figure 4:**
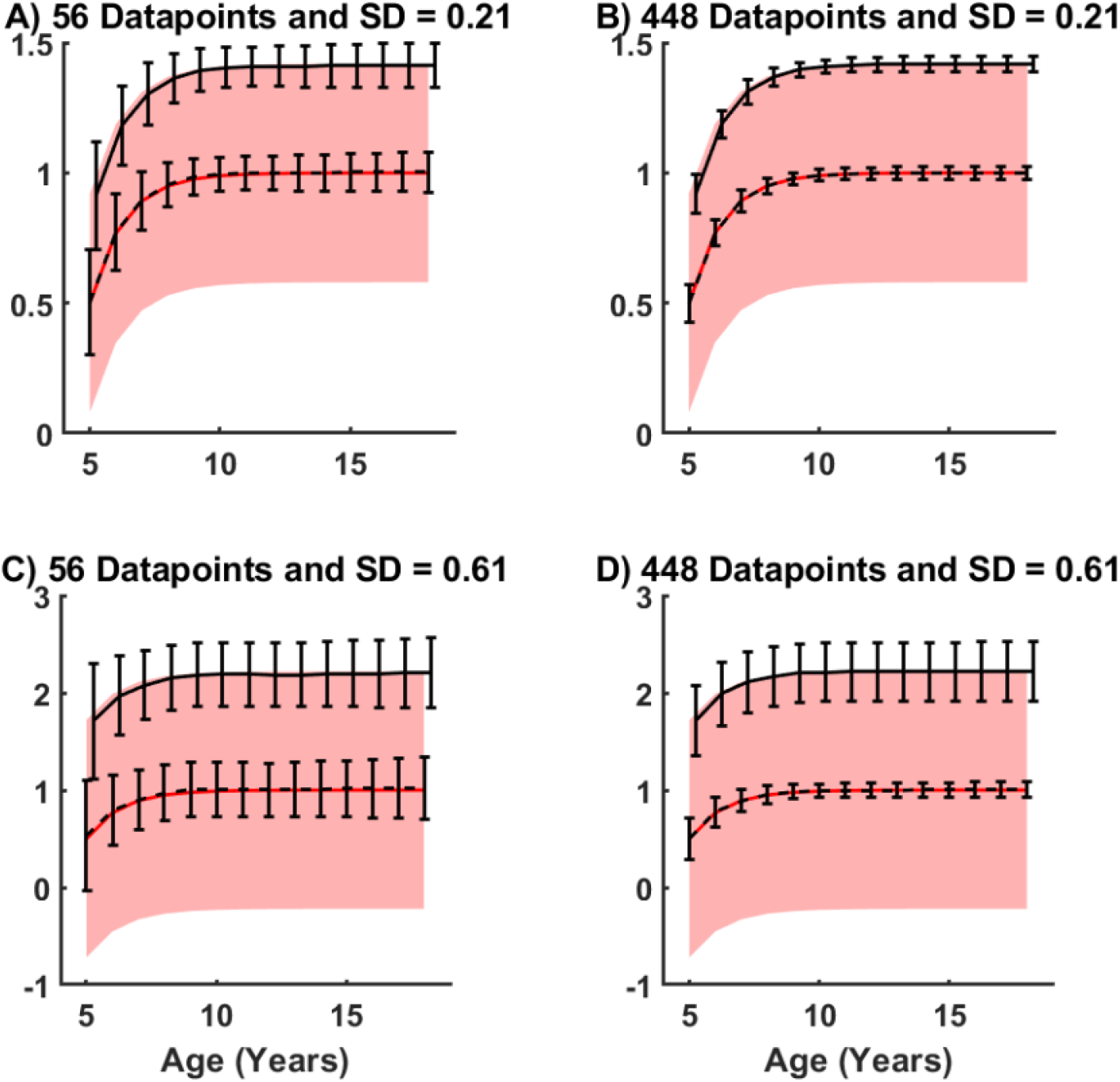
Input and output curve distributions for the growth to age 8 “near-adult” performance curve with two different input dataset sizes and SDs. The red line is the input curve, while the red region represents two input standard deviations above and below the input curve (95% of the input data lies within these bounds). The dashed line and error bars associated with it represent the mean and 95% CI of the output curve across all 10,000 simulations. The solid line and its associated error bars represent two output standard deviations above the mean output curve and its 95% CI across all 10,000 simulations. The 95% CI for the output standard deviation was calculated by adding the variance in the mean curve and output standard deviation. The error bars were offset on the age to clarify plotting where there was overlap between CIs. Only the error bars representing two output standard deviations above the mean output curve was plotted, as the two standard deviations below the mean would be a mirror image across the mean output curve.

On average, both the mean and 2SD closely matched the values for the input curves (<0.7% error), although the individual estimates display considerable error when compared to the input curve. At age 5, the 95% CI of the curve fits ranged from 0.3 to 0.7 (range = 0.4) even though the correct value was 0.5. At age 11, this range reduced to 0.12, and then increased slightly to 0.14 for age 18. The true 2SD value is 0.92 (0.5+2*0.21SD) at age 5 for this dataset and the estimated values averaged to 0.91; however, CI for 2SD varied substantially. In Fig. 4A, one can observe the range of estimates for 2SD (top curve) is 0.4 (0.72-1.12) which then drops to 0.15 at age 11 and increases slightly to 0.17 at age 18. Similar patterns of variability in the mean and SD are observed when SD and the data sample size are modified (Fig. 4B-D). For example, Fig. 4B highlights that the 95% CI for the mean and 2SD ranges reduce to 0.04 and 0.05, respectively at age 11.

Age differences around the mean fitted curve at the 95% CIs and dataset size together impact whether an individual child is identified as within the range of typical performance for that age. For two different values of SD, a plot of CIs for each dataset size vs age is displayed in Fig. 5A and 5B. The data in the figure represent the range of the 95% CI in units of input SD, i.e. a value of four for an input SD of 0.21 would mean the 95% CI has a range of 4*0.21 (0.84) in the original units. The units of input SD were chosen to facilitate comparisons across different input SDs. Fig. 5A shows the range of the 95% CIs at each age for all dataset sizes with an input SD of 0.21. Each line represents a different dataset size (shown beside the line) and the line thickness increases with dataset size. The 95% CI is largest at age 5, regardless of dataset size, and the CI range decreases with increasing age, with a slight increase at age 18. For example, in the dataset size of 112, the 95% CI range starts at 0.98 input SDs (0.98*0.21 = 0.206) at age 5, decreases to 0.30 input SDs (0.30*0.21 = 0.06) at age 11, and increases to 0.35 input SDs (0.35*0.21 = 0.07). The range of all 95% CIs decrease and become flatter, regardless of age, as dataset size increases. Fig. 5B with a SD of 0.61 illustrates that the same trends seen in Fig. 5A occur for different input SDs. For different input SDs, the range of the 95% CI for the same age and dataset size is affected in smaller dataset sizes, but not in larger datasets. For example, with the input SD of 0.61 and 112 datapoints, the 95% CI ranges are 0.95, 0.31, and 0.40, at age 5, 11 and 18 respectively. While the ranges differ when converted into the original units (0.95*0.61 = 0.58, 0.31*0.61 = 0.19, and 0.40*0.61 = 0.24 compared to 0.206, 0.06, and 0.07), the ranges are similar relative to the input SD.

**Figure 5:**
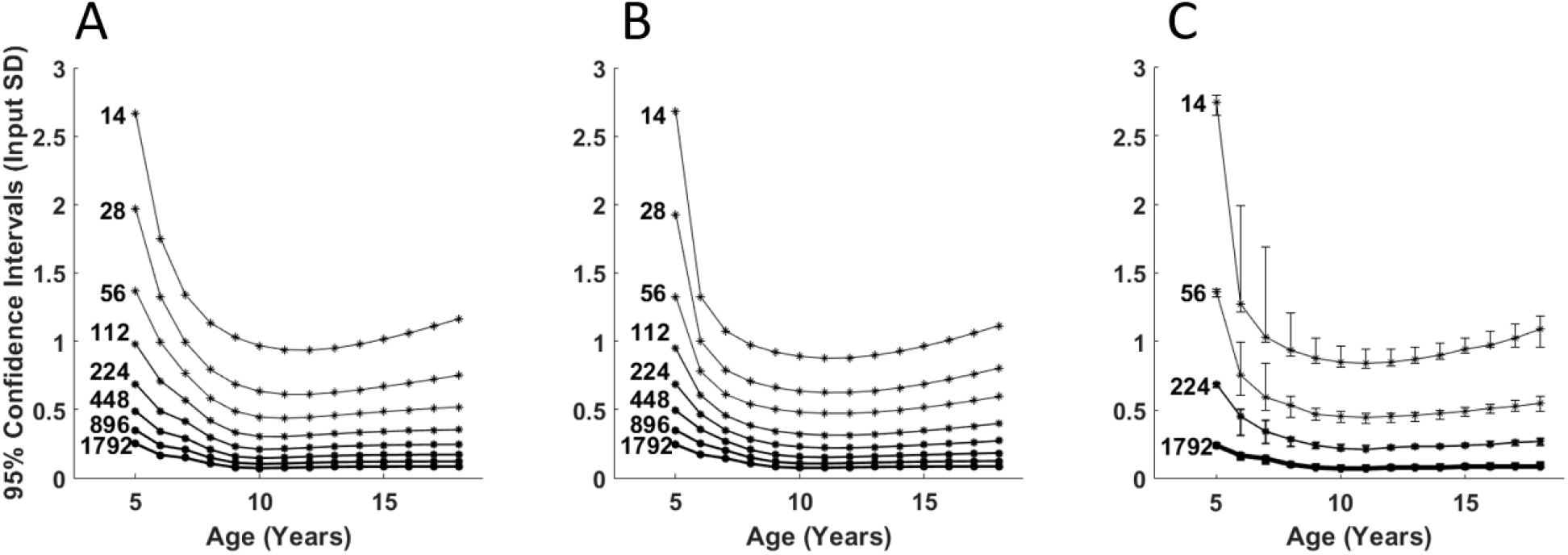
95% CIs of difference between input and output curves for each dataset size, plotted against age for two different input SDs, and span of the 95% CIs across all input SDs for the growth to age 8 input curve. Each line represents a different dataset size, with increasing line thickness corresponding to increasing dataset size. The corresponding dataset size is also shown to the left of each line. A) shows the results for an input SD of 0.21, and B) shows the results from an input SD of 0.61. As was noted in Fig. 4, the worst repeatability was at 5-years-old in all cases. This figure more clearly shows that the 95% CI reaches a minimum near the middle of the age range, then increases towards age 18. The increase is larger for smaller datasets. C) shows the span of the 95% CIs across all input SDs for the growth to age 8 input curve. The error bars represent the 2.5 and 97.5 percentiles of the calculated 95% CIs across all input SDs. The CI spans are largest for small dataset sizes, where accuracy and repeatability of the results are already poorest and decrease with increasing dataset size. The largest CI spans for the datasets with 14, 56, 224, and 1,792 datapoints are 0.77, 0.38, 0.19, and 0.04 SD respectively. Additionally, the CI spans change size at each age for all datasets, indicating there is variation between how well the curves are predicted at each age when the input SD is varied. The units for the plots are input standard deviation (Input SD).

The impact of all the different input SDs on the 95% CIs is shown in Fig. 5C. The 95% CIs shown in Fig. 5A and B were calculated for each input SD, then the distribution of these CIs was calculated and represented by the error bars. This distribution represented by the error bars will be called the CI span to avoid confusion. For clarity of the image, only four dataset sizes are shown: 14, 56, 224, and 1,792 datapoints. Smaller datasets show more variation in the CI spans for different input SDs and the variation decreases with increasing dataset size. The largest CI span for the datasets with 14, 56, 224, and 1,792 were 0.77, 0.38, 0.19, and 0.04 SDs, respectively. Although the repeatability of the curve fitting was always poorest for the 5-year-old datapoints, the largest variability among input SDs was typically at ages 6 or 7.

The same trends were found for all input curves (see Supplementary Material).

### Simulated Data with Non-Uniform Distributions

The same simulations were run with the age distribution of the participants who completed robotic assessments (Fig. 1). The simulations were run with the eight input curves, 15 SDs, but only two dataset sizes. The datasets had 288 and 576 datapoints to reflect the number of datapoints recorded for bimanual and unimanual tasks, respectively. The participant ages were not uniformly distributed across the age range (mean = 13.4, SD = 3.13, 190 male, 98 female). Most participants were between ages 11 and 17 with few participants under age 8. There were also nearly twice as many male participants as female participants.

Fig. 6 shows the same information as Fig. 5, but for the non-uniform dataset simulations with the decaying to age 8 input curve. Unlike the uniform datasets, the non-uniform datasets show a larger difference between the 95% CIs for the two plotted input SDs. At age 5 for the input SD of 0.21 (Fig. 6A), the 95% CIs were 2.95 and 1.79 input SD for the 288 and 576 datapoints, respectively, compared to 3.11 and 2.10 input SD for the larger input SD (Fig. 6B). The differences in 95% CIs between the input SDs were minimal for all other ages (within 0.13 input SD).

**Figure 6:**
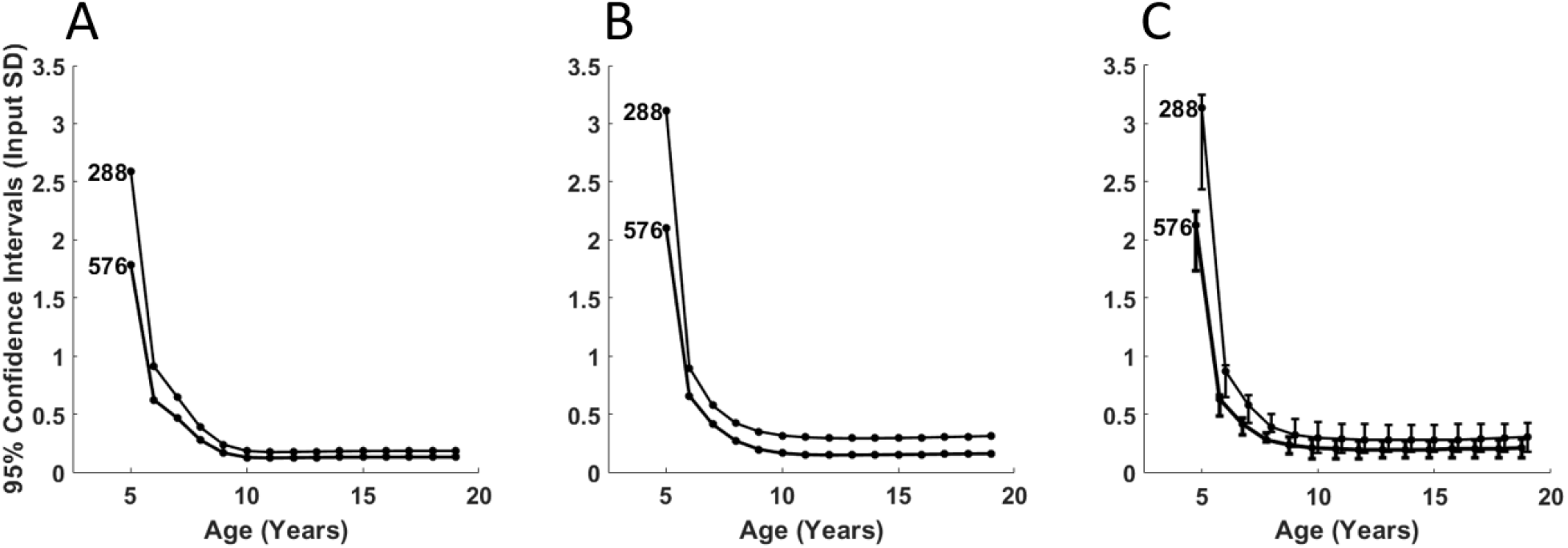
95% CIs of difference between input and output curves for each dataset size, plotted against age for two different input SDs for the non-uniform dataset, and span of the 95% CIs across all input SDs for the decaying to age 8 input curve and non-uniform dataset. Each line represents a different dataset size, with increasing line thickness corresponding to increasing dataset size. The corresponding dataset size is also shown to the left of each line. A) shows the results for an input SD of 0.21, and B) shows the results from an input SD of 0.61. C) shows the span of the 95% CIs across all input SDs for the decaying to age 8 input curve and non-uniform dataset. The error bars represent the 2.5 and 97.5 percentiles of the calculated 95% CIs across all input SDs. The CI spans are largest for small dataset sizes, where accuracy and repeatability of the results are already poorest and decrease with increasing dataset size. The largest CI spans for the datasets with 288 and 576 datapoints are 0.81 and 0.51 SD respectively. Additionally, the CI spans change size at each age for all datasets, indicating there is variation between how well the curves are predicted at each age when the input SD is varied. The units for the plots are input standard deviation (Input SD).

Fig. 6C shows the same information as Fig. 5C, but for the non-uniform dataset. The larger differences in 95% CIs among input SDs resulted in wider CI spans of the input SDs at age 5, but similar spans for the other ages when comparing the uniform and non-uniform datasets. The largest CI span for the 288 and 576 datapoint datasets were 0.81 and 0.51 SD, respectively. The CI spans for the non-uniform datasets are most comparable to the spans for the uniform datasets with 14 datapoints and 56 datapoints. The largest CI spans for the 14 and 56 datapoints were 0.77 and 0.38, respectively. The average CI spans at each age within the uniform datasets with 14 and 56 datapoints were 0.28 and 0.19, compared to 0.24 and 0.15 for the non-uniform 288 and 576 datapoint datasets.

To evaluate differences, the 95% CIs for uniform datasets and the closest (relative to size) non-uniform datasets at each age are plotted in Fig. 7. The comparisons were made with the input SD of 0.21. There were five participants who were five years old, so the 95% CIs of the dotted line at age 5 are from the uniform dataset with 56 datapoints, as there would be four datapoints at age 5. Fig. 7A shows the results from the 288 datapoint simulations, whereas Fig. 7B shows the results for the 576 datapoints. The 95% confidence intervals on the curve fitting differed when calculated from the simulations with a uniform vs non-uniform dataset. These differences were more evident in the younger ages.

**Figure 7:**
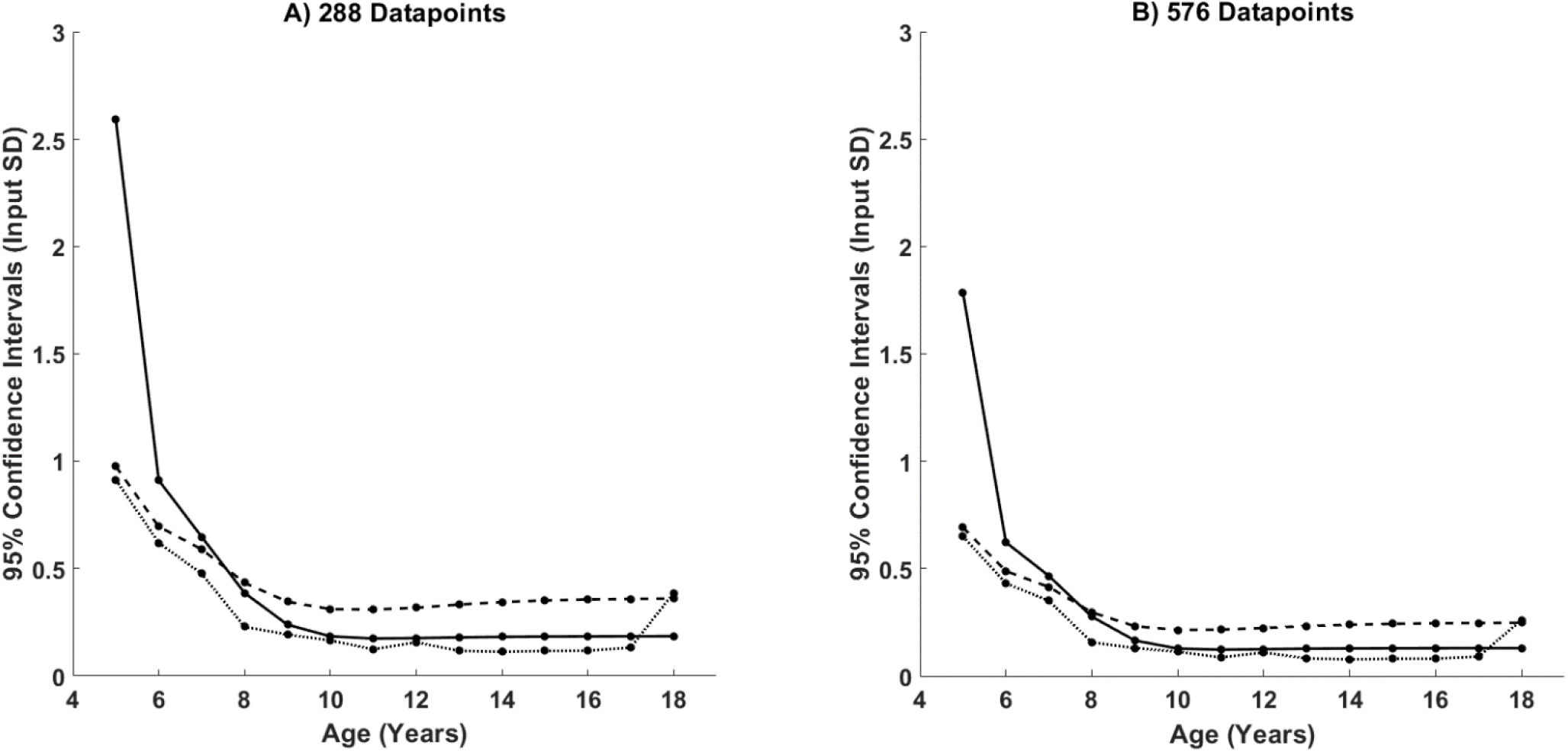
Comparisons of the 95% confidence intervals for the uniform and non-uniform input datasets with an input SD of 0.21. The solid line represents the results from the non-uniform input datasets, the dashed line represents the results from the closest uniform dataset (224 and 448 datapoints), and the dotted line represents the results of a uniform dataset with the closest number of participants at each age as the non-uniform dataset. The uniform dataset with the same number of datapoints at each age as the non-uniform dataset closely matched the results of the non-uniform dataset for ages 9 and up. The difference in the 95% CIs averaged 0.18 and 0.12 input SD for the 288 and 576 datapoint datasets, respectively.

### Assessing Accuracy and Repeatability of Curve Fitting: Fitting Confidence Intervals to Participant Data

The confidence intervals created by the simulations can be applied to curves that are fit to data collected from participants. These confidence intervals can be used to quantify the certainty of the identification of a deficit in a motor function performance measure. Curves were fit to Reaction Time (RT) collected from the Visually Guided Reaching (VGR) task. Fig. 8 shows the 95% confidence intervals on the curve fit to RT using the uniform and non-uniform datasets. Fig. 8A shows the 95% confidence intervals from the uniform datasets, whereas Fig. 8B shows the intervals from the non-uniform datasets. The difference between the 95% confidence interval widths from uniform and non-uniform datasets is 0.07 seconds at age 5, and decreases to 0.03 seconds at age 6. The differences are then below 0.01 seconds for all other ages, with the smallest difference, 0.0006 seconds, at age 16. These differences in the 95% confidence interval width range from 0.21% to 12% of the fitted curve’s value at each age.

**Figure 8:**
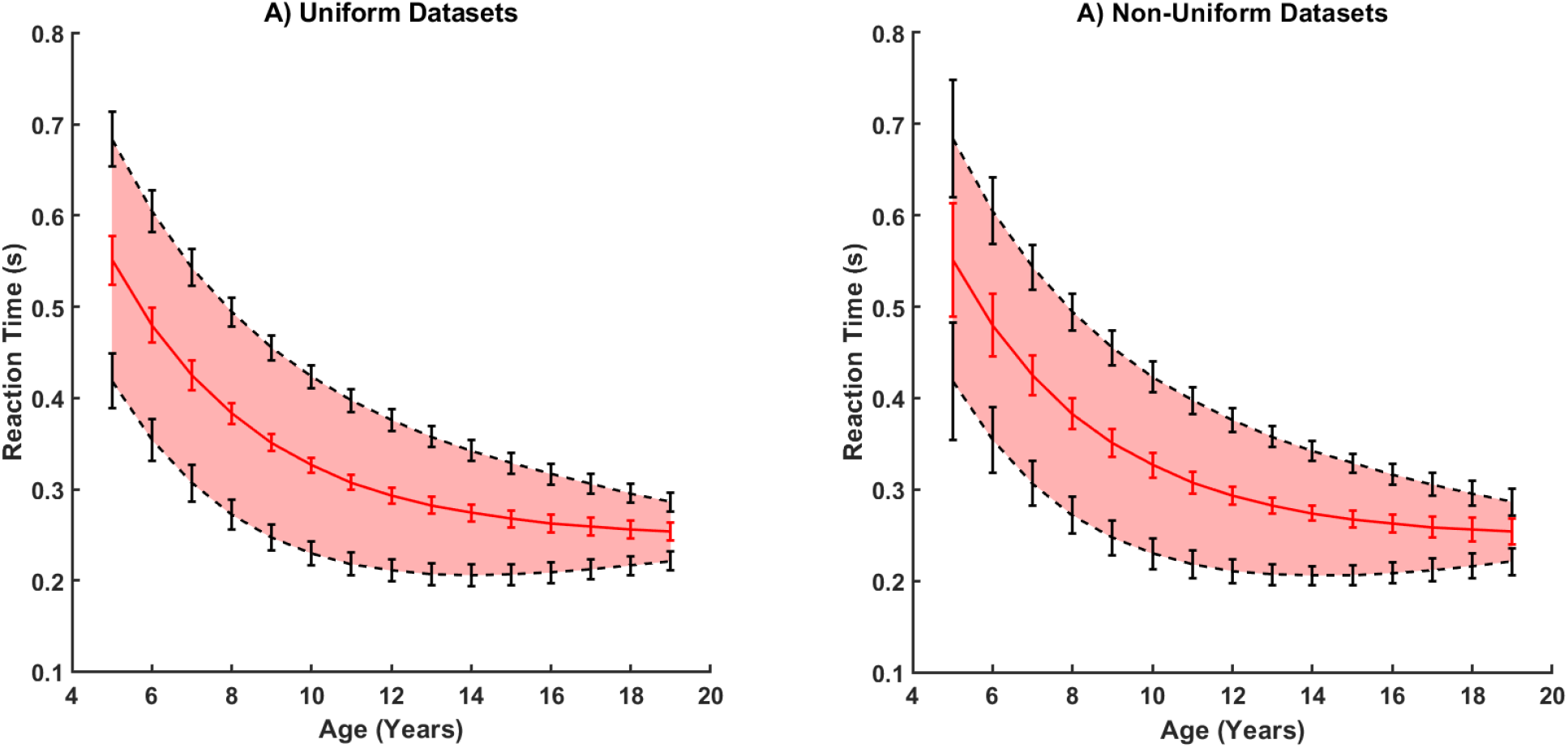
Comparisons of 95% confidence intervals using simulations with uniform and non-uniform datasets for the performance parameter RT. The red lines and shaded areas represent the fitted mean curve and +/-2SD from the mean. The confidence intervals on the fitted curve at age 5 are 0.06 seconds and 0.13 seconds for the uniform and non-uniform datasets, respectively. The difference in confidence interval width ranges from 0.0006 to 0.07. The difference drops from 0.07 at age 5 to 0.03 at age 6, and is smallest at age 16.

From ages 9-18, the average difference between the 95% CIs generated from the uniform and non-uniform datasets with matched numbers of participants were 0.18 and 0.12 input SD for the 288 and 576 datapoint datasets, respectively.

## Discussion

An algorithm for creating non-linear developmental models of motor performance that accounted for changing variability was created. The repeatability of the curve fitting was measured using simulated data with both a uniform age distribution and a non-uniform distribution similar to the participant data. The simulations tested how dataset size, variability in the data, and curvature of the data affected the curve fitting.

### Repeatability of Curve Fitting

We assessed and presented curve fitting performance rather than setting desired goals for accuracy to facilitate the use of the presented method and findings to inform future work (see Supplementary Material). When collecting data for clinical studies, the number of participants is often difficult to determine a priori; however, the current findings could be utilized to obtain insight about the limitations and benefits of different sized groups and to inform power analyses. Similarly, the information presented here could be used to determine the participant numbers needed to enable curve fitting that is sufficiently accurate for experimental purposes.

The simulations with different sample sizes, variability in data, and underlying curvature of the data highlighted that the sample size tends to have the largest impact on accuracy and repeatability for these non-linear regressions for both uniform and non-uniform data distributions. Differences in accuracy and repeatability among different input standard deviations and input curves decreased with increasing sample size. These results indicate that any future work assessing the accuracy and repeatability of non-linear curve fitting may not need to assess different input SDs and curves, but simply use one SD and input curve and vary the sample size to determine general accuracy and repeatability of the curve fitting.

The repeatability of the curve fitting improved in a decaying exponential fashion with increasing dataset size. The average 95% CIs across all input SDs at the beginning of the curve decreased from 2.7 to 0.24 input SD, a ten-fold decrease, from 14 to 1,792 datapoints respectively for the input curve representing growth to age 8 to reach “near-adult” performance. Most of the decrease was observed for datasets smaller than 224. Once the dataset size reached 224 datapoints, the average 95% CI was 0.68, so 82% of the overall improvement occurred between sample sizes from 14 to 224 datapoint.

The smaller improvements in repeatability above a dataset of 224 datapoints are consistent with another study on logistic regression for modelling complications from radiotherapy [21]. The authors found that improvements in predictive power of the models were marginal among datasets with more than 200 datapoints but were larger among smaller datasets [21]. Schönbrodt and Perugini determined stabilized correlations occurred at 250 datapoints which is similar to our 224 datapoints [22]. These results indicate that using non-linear models to create typical performance ranges requires above 200 datapoints to create ranges which could accurately differentiate typical from atypical performance. More simulations would be necessary to determine an exact number of datapoints required.

Regardless of input dataset size, SD, or type of input curve, the poorest accuracy and repeatability was always found at the beginning of the curve (age 5). These results indicate that it may be useful to either use weighting methods in curve fitting to improve accuracy at the beginning of the curve, or to collect more datapoints at the boundaries of the curve. When using this approach, (i.e. weighting the curve fitting or increasing the number of datapoints at the beginning of the curve) caution must be taken to accurately fit that portion of the curve. For example, if there are more datapoints at age 5, then minimizing the error on those datapoints will be more important than minimizing error where there are fewer datapoints. However, an increased focus on minimizing error at the start of the curve may decrease the accuracy of the fit at the end of the curve, so care would need to be taken to ensure a sufficiently accurate and repeatable fit along the entire curve.

At the beginning of the curve, where collected data were most sparse, the repeatability of the curve fitting was the lowest and it was least comparable to the results from a uniform dataset. The 95% confidence intervals on the curve fitting were similar between the non-uniform and uniform datasets from ages 9 and up (within 0.18 input SD). These results indicate that the simulations from the uniform dataset may be suitable to create 95% confidence intervals on new fitted curves for ages 9 and up for this particular dataset. The poor repeatability from ages 5 to 8 indicate that using these normative models to identify motor deficits in children that age may not be appropriate.

The curve fitting algorithm has been shown to be a reliable tool for fitting non-linear functions to simulated data and determining distributions of the input data. The algorithm could prove to be a useful tool for creating future models of experimental data with known accuracy and repeatability based on dataset size and data variability. If 95% CIs are required for similar simulations, researchers may only need to assess one input curve using multiple dataset sizes, as the dataset size is the most important factor in the repeatability of the curve fitting.

### Confidence Intervals Based on Normative Models

Normative models were created for the Reaction Time (RT) data collected from the Visually Guided Reaching (VGR) task. The confidence intervals generated from the uniformly distributed and non-uniformly distributed datasets were compared across the fitted curve to the RT data. The confidence intervals were similar from ages 9-18 and the simulations from the uniform distributions could be used for future normative models, even if they are generated from non-uniformly distributed data. A more thorough analysis of the impact of non-uniformity of input data distributions on curve fitting should be considered.

The authors have not found previous studies that included confidence intervals in normative models of motor development in children. Including confidence intervals quantifies the reliability of the curve fitting and improves accuracy in the identification of motor function deficits. Reporting confidence intervals on statistical tests is common in other disciplines and should be more common for these types of normative models to allow for a full understanding of the strength of the conclusions drawn from the results.

### Limitations

The curve fitting algorithm was given a very good initial estimate of the coefficients used to generate the data (within 1% of each coefficient), and the accuracy and repeatability were not compared for different ranges of input coefficients. Poor initial estimates would most likely result in poorer fits to the data. The issue of poorer fits can be partially fixed by changing parts of the curve fitting function, such as the number of optimization iterations or optimization parameters. These additional solutions have not been explored in this work.

### Clinical Implications

The results of the simulations can be applied to normative models of motor development to inform the certainty of identification of a deficit. Clinicians can use normative models to identify impairments and target and track therapies. The inclusion of confidence intervals on these models will help clinicians decide if someone has a deficit, or if they require more assessment to make an accurate decision.

Simulated data were used to assess how dataset size, curvature in the data, and variability in the data affect the accuracy and repeatability of the curve fitting. The simulations for curve fitting were completed to fill the gap in which there was a lack in reporting of true confidence in normative ranges of motor development in children. The curvature and variation of the input data did not affect the confidence intervals in a meaningful way; only the sample size had a large effect on the confidence intervals. The accuracy and repeatability were proportional to the variability in the data; however, when normalized to the variability, the accuracy and repeatability were similar among the different curve fits. Increasing dataset size improved the accuracy and repeatability of the curve fitting. Most of the improvement from the uniformly distributed data was observed at the dataset size of 224 datapoints. The results of these simulations can be used to determine a necessary sample size for a desired confidence in a curve fit, or to assess the confidence in a curve fit based on the collected dataset size. The identification of the important parameters to explore when performing these types of simulations may encourage other researchers to complete these simulations and report confidence intervals in their results.

The results from the simulations also showed that the confidence intervals of an exponential fit are not constant across the entire range of the input variable, age in this case. These results highlight that using a single value for confidence interval width may be inappropriate for this type of analysis. Confidence intervals should be generated for different input values, rather than using one value across the entire range of input values.

## Conflict of Interest

S.H. Scott is the Chief Scientific Officer and co-founder of Kinarm, the company that develops the robot and software used in this article.

## Acknowledgements

The authors would like to thank: S. Appaqaq, H. Bretzke, E. Heming, and K. Moore, for their technical support; and E. Aleska, E. Castillo, J. Empey, A. Gowthorpe, E. Hoskin, L. Jansen, E. Johannessen, S. Klinger, A. Lax-Vanek, A. Lopez, L. Munro, E. Neff, E. Perfect, E. Rendall, O. Roud, N. Smith, R. Spender, and K. Van-Til for their help with data collection. The authors would also like to thank C. Lowrey for her help with editing the manuscript.

