## Supplement tables and figures. for "Quantifying Effects of Dataset Size, Data Variability, and Data Curvature on Modelling of Simulated Age-Related Motor Development Data"

### Supplementary Material

This document contains supplementary material for the article, “Quantifying Effects of Dataset Size, Data Variability, and Data Curvature on Modelling of Simulated Age-Related Motor Development Data”. The document contains tables of the CI spans (span of the confidence interval, CI), and figures of the CI spans for each input curve described in the article.

The tables show the data for the CI spans for each input curve at each age for all input dataset sizes. This is the information used to create Figs. 5 and 6 in the article, and the figures included in this document. From these tables, the accuracy and repeatability of the fitted curve at different ages for different sample sizes can be determined. The data in the table are written as mean performance (bottom of CI span, top of CI span), and are in units of input standard deviation (SD).

### Uniform Datasets

#### Growth Curves

##### Growth to age 8

Table 1: CI spans for the growth to age 8 input curve.

| Age | Dataset Size (number of datapoints) | | | | | | | |
|---|---|---|---|---|---|---|---|---|
| | 14 | 28 | 56 | 112 | 224 | 448 | 896 | 1,792 |
| 5 | 2.747 (2.651, 2.798) | 1.932 (1.889, 1.976) | 1.361 (1.326, 1.384) | 0.962 (0.935, 0.986) | 0.685 (0.665, 0.699) | 0.488 (0.466, 0.494) | 0.347 (0.331, 0.351) | 0.244 (0.235, 0.251) |
| 6 | 1.275 (1.217, 1.99) | 0.972 (0.847, 1.417) | 0.751 (0.608, 0.993) | 0.589 (0.428, 0.711) | 0.454 (0.315, 0.502) | 0.336 (0.24, 0.356) | 0.241 (0.187, 0.253) | 0.17 (0.145, 0.178) |
| 7 | 1.033 (0.993, 1.691) | 0.783 (0.687, 1.184) | 0.596 (0.496, 0.841) | 0.457 (0.353, 0.592) | 0.347 (0.255, 0.422) | 0.264 (0.193, 0.297) | 0.196 (0.149, 0.211) | 0.144 (0.115, 0.15) |
| 8 | 0.933 (0.892, 1.205) | 0.703 (0.623, 0.843) | 0.531 (0.447, 0.603) | 0.39 (0.322, 0.424) | 0.284 (0.232, 0.3) | 0.204 (0.173, 0.21) | 0.146 (0.133, 0.151) | 0.105 (0.098, 0.106) |
| 9 | 0.882 (0.839, 1.028) | 0.64 (0.591, 0.737) | 0.471 (0.421, 0.513) | 0.344 (0.305, 0.372) | 0.241 (0.219, 0.262) | 0.17 (0.155, 0.18) | 0.118 (0.112, 0.13) | 0.082 (0.078, 0.095) |
| 10 | 0.848 (0.813, 0.967) | 0.614 (0.576, 0.708) | 0.45 (0.407, 0.487) | 0.319 (0.289, 0.355) | 0.222 (0.205, 0.248) | 0.157 (0.144, 0.171) | 0.107 (0.102, 0.124) | 0.075 (0.071, 0.091) |
| 11 | 0.845 (0.808, 0.941) | 0.61 (0.574, 0.694) | 0.445 (0.403, 0.475) | 0.314 (0.294, 0.349) | 0.216 (0.211, 0.241) | 0.155 (0.15, 0.167) | 0.107 (0.106, 0.121) | 0.076 (0.074, 0.089) |
| 12 | 0.851 (0.818, 0.941) | 0.613 (0.581, 0.693) | 0.45 (0.407, 0.474) | 0.316 (0.296, 0.349) | 0.224 (0.212, 0.24) | 0.159 (0.154, 0.166) | 0.112 (0.109, 0.121) | 0.078 (0.077, 0.089) |
| 13 | 0.871 (0.841, 0.957) | 0.625 (0.597, 0.703) | 0.462 (0.417, 0.483) | 0.325 (0.303, 0.354) | 0.232 (0.217, 0.244) | 0.164 (0.159, 0.169) | 0.115 (0.113, 0.123) | 0.081 (0.079, 0.091) |
| 14 | 0.903 (0.874, 0.987) | 0.646 (0.62, 0.724) | 0.474 (0.432, 0.498) | 0.334 (0.313, 0.364) | 0.238 (0.224, 0.25) | 0.168 (0.164, 0.174) | 0.118 (0.117, 0.127) | 0.083 (0.081, 0.093) |
| 15 | 0.942 (0.912, 1.026) | 0.673 (0.648, 0.751) | 0.491 (0.452, 0.518) | 0.346 (0.327, 0.377) | 0.241 (0.233, 0.258) | 0.171 (0.169, 0.181) | 0.12 (0.119, 0.132) | 0.085 (0.083, 0.096) |
| 16 | 0.975 (0.955, 1.073) | 0.699 (0.676, 0.784) | 0.509 (0.475, 0.541) | 0.361 (0.34, 0.392) | 0.249 (0.242, 0.269) | 0.179 (0.171, 0.189) | 0.123 (0.12, 0.137) | 0.086 (0.084, 0.1) |
| 17 | 1.025 (0.958, 1.126) | 0.729 (0.678, 0.821) | 0.529 (0.489, 0.568) | 0.369 (0.341, 0.409) | 0.261 (0.242, 0.281) | 0.185 (0.171, 0.198) | 0.125 (0.12, 0.143) | 0.087 (0.084, 0.105) |
| 18 | 1.087 (0.959, 1.184) | 0.769 (0.679, 0.862) | 0.549 (0.491, 0.597) | 0.389 (0.341, 0.427) | 0.271 (0.243, 0.294) | 0.192 (0.172, 0.208) | 0.126 (0.12, 0.15) | 0.087 (0.084, 0.109) |

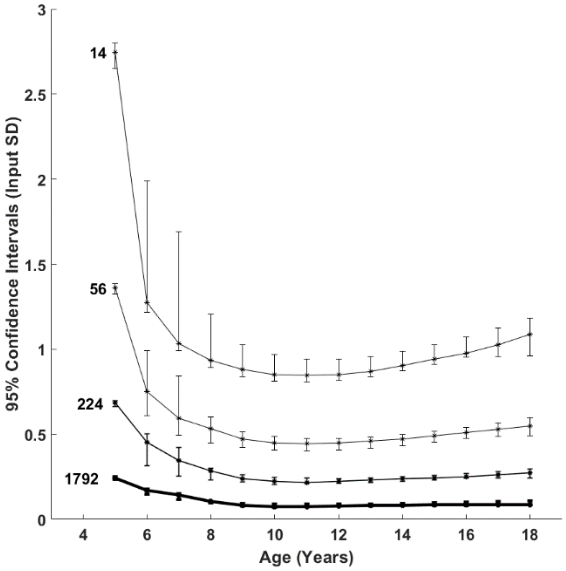


Fig. 1. Span of the 95% CIs across all input SDs for the growth to age 8 input curve. The error bars represent the 2.5 and 97.5 percentiles of the calculated 95% CIs across all input SDs.

#### Growth to age 10

Table 2: CI spans for the growth to age 10 input curve.

| Age | Dataset Size (number of datapoints) | | | | | | | |
|---|---|---|---|---|---|---|---|---|
| | 14 | 28 | 56 | 112 | 224 | 448 | 896 | 1,792 |
| 5 | 2.747 (2.651, 2.798) | 1.932 (1.889, 1.976) | 1.361 (1.326, 1.384) | 0.962 (0.935, 0.986) | 0.685 (0.665, 0.699) | 0.488 (0.466, 0.494) | 0.347 (0.331, 0.351) | 0.244 (0.235, 0.251) |
| 6 | 1.275 (1.217, 1.99) | 0.972 (0.847, 1.417) | 0.751 (0.608, 0.993) | 0.589 (0.428, 0.711) | 0.454 (0.315, 0.502) | 0.336 (0.24, 0.356) | 0.241 (0.187, 0.253) | 0.17 (0.145, 0.178) |
| 7 | 1.033 (0.993, 1.691) | 0.783 (0.687, 1.184) | 0.596 (0.496, 0.841) | 0.457 (0.353, 0.592) | 0.347 (0.255, 0.422) | 0.264 (0.193, 0.297) | 0.196 (0.149, 0.211) | 0.144 (0.115, 0.15) |
| 8 | 0.933 (0.892, 1.205) | 0.703 (0.623, 0.843) | 0.531 (0.447, 0.603) | 0.39 (0.322, 0.424) | 0.284 (0.232, 0.3) | 0.204 (0.173, 0.21) | 0.146 (0.133, 0.151) | 0.105 (0.098, 0.106) |
| 9 | 0.882 (0.839, 1.028) | 0.64 (0.591, 0.737) | 0.471 (0.421, 0.513) | 0.344 (0.305, 0.372) | 0.241 (0.219, 0.262) | 0.17 (0.155, 0.18) | 0.118 (0.112, 0.13) | 0.082 (0.078, 0.095) |
| 10 | 0.848 (0.813, 0.967) | 0.614 (0.576, 0.708) | 0.45 (0.407, 0.487) | 0.319 (0.289, 0.355) | 0.222 (0.205, 0.248) | 0.157 (0.144, 0.171) | 0.107 (0.102, 0.124) | 0.075 (0.071, 0.091) |
| 11 | 0.845 (0.808, 0.941) | 0.61 (0.574, 0.694) | 0.445 (0.403, 0.475) | 0.314 (0.294, 0.349) | 0.216 (0.211, 0.241) | 0.155 (0.15, 0.167) | 0.107 (0.106, 0.121) | 0.076 (0.074, 0.089) |
| 12 | 0.851 (0.818, 0.941) | 0.613 (0.581, 0.693) | 0.45 (0.407, 0.474) | 0.316 (0.296, 0.349) | 0.224 (0.212, 0.24) | 0.159 (0.154, 0.166) | 0.112 (0.109, 0.121) | 0.078 (0.077, 0.089) |
| 13 | 0.871 (0.841, 0.957) | 0.625 (0.597, 0.703) | 0.462 (0.417, 0.483) | 0.325 (0.303, 0.354) | 0.232 (0.217, 0.244) | 0.164 (0.159, 0.169) | 0.115 (0.113, 0.123) | 0.081 (0.079, 0.091) |
| 14 | 0.903 (0.874, 0.987) | 0.646 (0.62, 0.724) | 0.474 (0.432, 0.498) | 0.334 (0.313, 0.364) | 0.238 (0.224, 0.25) | 0.168 (0.164, 0.174) | 0.118 (0.117, 0.127) | 0.083 (0.081, 0.093) |
| 15 | 0.942 (0.912, 1.026) | 0.673 (0.648, 0.751) | 0.491 (0.452, 0.518) | 0.346 (0.327, 0.377) | 0.241 (0.233, 0.258) | 0.171 (0.169, 0.181) | 0.12 (0.119, 0.132) | 0.085 (0.083, 0.096) |
| 16 | 0.975 (0.955, 1.073) | 0.699 (0.676, 0.784) | 0.509 (0.475, 0.541) | 0.361 (0.34, 0.392) | 0.249 (0.242, 0.269) | 0.179 (0.171, 0.189) | 0.123 (0.12, 0.137) | 0.086 (0.084, 0.1) |
| 17 | 1.025 (0.958, 1.126) | 0.729 (0.678, 0.821) | 0.529 (0.489, 0.568) | 0.369 (0.341, 0.409) | 0.261 (0.242, 0.281) | 0.185 (0.171, 0.198) | 0.125 (0.12, 0.143) | 0.087 (0.084, 0.105) |
| 18 | 1.087 (0.959, 1.184) | 0.769 (0.679, 0.862) | 0.549 (0.491, 0.597) | 0.389 (0.341, 0.427) | 0.271 (0.243, 0.294) | 0.192 (0.172, 0.208) | 0.126 (0.12, 0.15) | 0.087 (0.084, 0.109) |

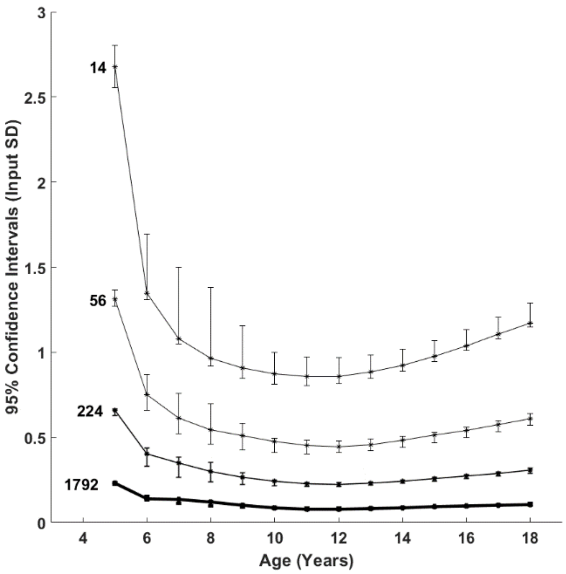


Fig. 2. Span of the 95% CIs across all input SDs for the growth to age 10 input curve. The error bars represent the 2.5 and 97.5 percentiles of the calculated 95% CIs across all input SDs.

#### Growth to age 12

Table 3: CI spans for the growth to age 12 input curve.

| Age | Dataset Size (number of datapoints) | | | | | | | |
|---|---|---|---|---|---|---|---|---|
| | 14 | 28 | 56 | 112 | 224 | 448 | 896 | 1,792 |
| 5 | 2.747 (2.651, 2.798) | 1.932 (1.889, 1.976) | 1.361 (1.326, 1.384) | 0.962 (0.935, 0.986) | 0.685 (0.665, 0.699) | 0.488 (0.466, 0.494) | 0.347 (0.331, 0.351) | 0.244 (0.235, 0.251) |
| 6 | 1.275 (1.217, 1.99) | 0.972 (0.847, 1.417) | 0.751 (0.608, 0.993) | 0.589 (0.428, 0.711) | 0.454 (0.315, 0.502) | 0.336 (0.24, 0.356) | 0.241 (0.187, 0.253) | 0.17 (0.145, 0.178) |
| 7 | 1.033 (0.993, 1.691) | 0.783 (0.687, 1.184) | 0.596 (0.496, 0.841) | 0.457 (0.353, 0.592) | 0.347 (0.255, 0.422) | 0.264 (0.193, 0.297) | 0.196 (0.149, 0.211) | 0.144 (0.115, 0.15) |
| 8 | 0.933 (0.892, 1.205) | 0.703 (0.623, 0.843) | 0.531 (0.447, 0.603) | 0.39 (0.322, 0.424) | 0.284 (0.232, 0.3) | 0.204 (0.173, 0.21) | 0.146 (0.133, 0.151) | 0.105 (0.098, 0.106) |
| 9 | 0.882 (0.839, 1.028) | 0.64 (0.591, 0.737) | 0.471 (0.421, 0.513) | 0.344 (0.305, 0.372) | 0.241 (0.219, 0.262) | 0.17 (0.155, 0.18) | 0.118 (0.112, 0.13) | 0.082 (0.078, 0.095) |
| 10 | 0.848 (0.813, 0.967) | 0.614 (0.576, 0.708) | 0.45 (0.407, 0.487) | 0.319 (0.289, 0.355) | 0.222 (0.205, 0.248) | 0.157 (0.144, 0.171) | 0.107 (0.102, 0.124) | 0.075 (0.071, 0.091) |
| 11 | 0.845 (0.808, 0.941) | 0.61 (0.574, 0.694) | 0.445 (0.403, 0.475) | 0.314 (0.294, 0.349) | 0.216 (0.211, 0.241) | 0.155 (0.15, 0.167) | 0.107 (0.106, 0.121) | 0.076 (0.074, 0.089) |
| 12 | 0.851 (0.818, 0.941) | 0.613 (0.581, 0.693) | 0.45 (0.407, 0.474) | 0.316 (0.296, 0.349) | 0.224 (0.212, 0.24) | 0.159 (0.154, 0.166) | 0.112 (0.109, 0.121) | 0.078 (0.077, 0.089) |
| 13 | 0.871 (0.841, 0.957) | 0.625 (0.597, 0.703) | 0.462 (0.417, 0.483) | 0.325 (0.303, 0.354) | 0.232 (0.217, 0.244) | 0.164 (0.159, 0.169) | 0.115 (0.113, 0.123) | 0.081 (0.079, 0.091) |
| 14 | 0.903 (0.874, 0.987) | 0.646 (0.62, 0.724) | 0.474 (0.432, 0.498) | 0.334 (0.313, 0.364) | 0.238 (0.224, 0.25) | 0.168 (0.164, 0.174) | 0.118 (0.117, 0.127) | 0.083 (0.081, 0.093) |
| 15 | 0.942 (0.912, 1.026) | 0.673 (0.648, 0.751) | 0.491 (0.452, 0.518) | 0.346 (0.327, 0.377) | 0.241 (0.233, 0.258) | 0.171 (0.169, 0.181) | 0.12 (0.119, 0.132) | 0.085 (0.083, 0.096) |
| 16 | 0.975 (0.955, 1.073) | 0.699 (0.676, 0.784) | 0.509 (0.475, 0.541) | 0.361 (0.34, 0.392) | 0.249 (0.242, 0.269) | 0.179 (0.171, 0.189) | 0.123 (0.12, 0.137) | 0.086 (0.084, 0.1) |
| 17 | 1.025 (0.958, 1.126) | 0.729 (0.678, 0.821) | 0.529 (0.489, 0.568) | 0.369 (0.341, 0.409) | 0.261 (0.242, 0.281) | 0.185 (0.171, 0.198) | 0.125 (0.12, 0.143) | 0.087 (0.084, 0.105) |
| 18 | 1.087 (0.959, 1.184) | 0.769 (0.679, 0.862) | 0.549 (0.491, 0.597) | 0.389 (0.341, 0.427) | 0.271 (0.243, 0.294) | 0.192 (0.172, 0.208) | 0.126 (0.12, 0.15) | 0.087 (0.084, 0.109) |

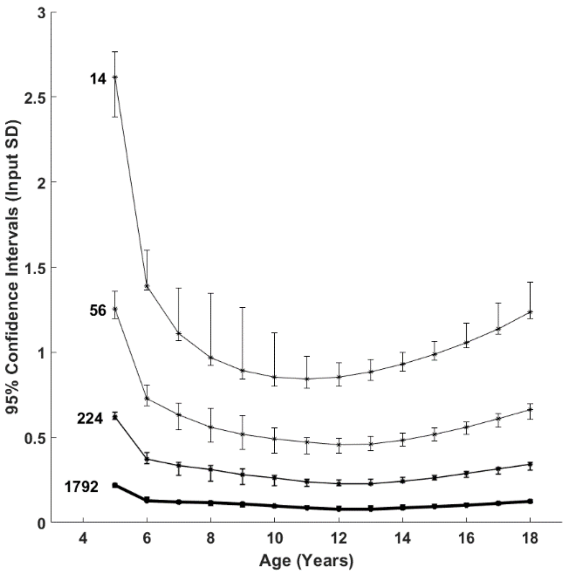


Fig. 3. Span of the 95% CIs across all input SDs for the growth to age 12 input curve. The error bars represent the 2.5 and 97.5 percentiles of the calculated 95% CIs across all input SDs.

#### Growth to age 14

Table 4: CI spans for the growth to age 14 input curve.

| Age | Dataset Size (number of datapoints) | | | | | | | |
|---|---|---|---|---|---|---|---|---|
| | 14 | 28 | 56 | 112 | 224 | 448 | 896 | 1,792 |
| 5 | 2.747 (2.651, 2.798) | 1.932 (1.889, 1.976) | 1.361 (1.326, 1.384) | 0.962 (0.935, 0.986) | 0.685 (0.665, 0.699) | 0.488 (0.466, 0.494) | 0.347 (0.331, 0.351) | 0.244 (0.235, 0.251) |
| 6 | 1.275 (1.217, 1.99) | 0.972 (0.847, 1.417) | 0.751 (0.608, 0.993) | 0.589 (0.428, 0.711) | 0.454 (0.315, 0.502) | 0.336 (0.24, 0.356) | 0.241 (0.187, 0.253) | 0.17 (0.145, 0.178) |
| 7 | 1.033 (0.993, 1.691) | 0.783 (0.687, 1.184) | 0.596 (0.496, 0.841) | 0.457 (0.353, 0.592) | 0.347 (0.255, 0.422) | 0.264 (0.193, 0.297) | 0.196 (0.149, 0.211) | 0.144 (0.115, 0.15) |
| 8 | 0.933 (0.892, 1.205) | 0.703 (0.623, 0.843) | 0.531 (0.447, 0.603) | 0.39 (0.322, 0.424) | 0.284 (0.232, 0.3) | 0.204 (0.173, 0.21) | 0.146 (0.133, 0.151) | 0.105 (0.098, 0.106) |
| 9 | 0.882 (0.839, 1.028) | 0.64 (0.591, 0.737) | 0.471 (0.421, 0.513) | 0.344 (0.305, 0.372) | 0.241 (0.219, 0.262) | 0.17 (0.155, 0.18) | 0.118 (0.112, 0.13) | 0.082 (0.078, 0.095) |
| 10 | 0.848 (0.813, 0.967) | 0.614 (0.576, 0.708) | 0.45 (0.407, 0.487) | 0.319 (0.289, 0.355) | 0.222 (0.205, 0.248) | 0.157 (0.144, 0.171) | 0.107 (0.102, 0.124) | 0.075 (0.071, 0.091) |
| 11 | 0.845 (0.808, 0.941) | 0.61 (0.574, 0.694) | 0.445 (0.403, 0.475) | 0.314 (0.294, 0.349) | 0.216 (0.211, 0.241) | 0.155 (0.15, 0.167) | 0.107 (0.106, 0.121) | 0.076 (0.074, 0.089) |
| 12 | 0.851 (0.818, 0.941) | 0.613 (0.581, 0.693) | 0.45 (0.407, 0.474) | 0.316 (0.296, 0.349) | 0.224 (0.212, 0.24) | 0.159 (0.154, 0.166) | 0.112 (0.109, 0.121) | 0.078 (0.077, 0.089) |
| 13 | 0.871 (0.841, 0.957) | 0.625 (0.597, 0.703) | 0.462 (0.417, 0.483) | 0.325 (0.303, 0.354) | 0.232 (0.217, 0.244) | 0.164 (0.159, 0.169) | 0.115 (0.113, 0.123) | 0.081 (0.079, 0.091) |
| 14 | 0.903 (0.874, 0.987) | 0.646 (0.62, 0.724) | 0.474 (0.432, 0.498) | 0.334 (0.313, 0.364) | 0.238 (0.224, 0.25) | 0.168 (0.164, 0.174) | 0.118 (0.117, 0.127) | 0.083 (0.081, 0.093) |
| 15 | 0.942 (0.912, 1.026) | 0.673 (0.648, 0.751) | 0.491 (0.452, 0.518) | 0.346 (0.327, 0.377) | 0.241 (0.233, 0.258) | 0.171 (0.169, 0.181) | 0.12 (0.119, 0.132) | 0.085 (0.083, 0.096) |
| 16 | 0.975 (0.955, 1.073) | 0.699 (0.676, 0.784) | 0.509 (0.475, 0.541) | 0.361 (0.34, 0.392) | 0.249 (0.242, 0.269) | 0.179 (0.171, 0.189) | 0.123 (0.12, 0.137) | 0.086 (0.084, 0.1) |
| 17 | 1.025 (0.958, 1.126) | 0.729 (0.678, 0.821) | 0.529 (0.489, 0.568) | 0.369 (0.341, 0.409) | 0.261 (0.242, 0.281) | 0.185 (0.171, 0.198) | 0.125 (0.12, 0.143) | 0.087 (0.084, 0.105) |
| 18 | 1.087 (0.959, 1.184) | 0.769 (0.679, 0.862) | 0.549 (0.491, 0.597) | 0.389 (0.341, 0.427) | 0.271 (0.243, 0.294) | 0.192 (0.172, 0.208) | 0.126 (0.12, 0.15) | 0.087 (0.084, 0.109) |

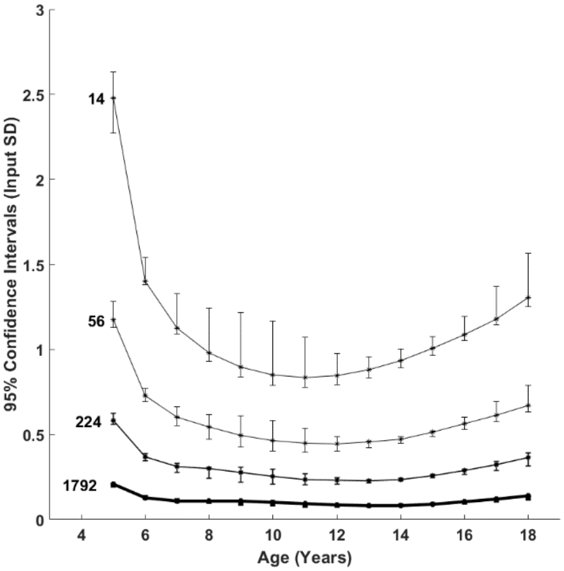


Fig. 4. Span of the 95% CIs across all input SDs for the growth to age 14 input curve. The error bars represent the 2.5 and 97.5 percentiles of the calculated 95% CIs across all input SDs.

### Decay Curves

#### Decay to age 8

Table 5: CI spans for the decay to age 8 input curve.

| Age | Dataset Size (number of datapoints) | | | | | | | |
|---|---|---|---|---|---|---|---|---|
| | 14 | 28 | 56 | 112 | 224 | 448 | 896 | 1,792 |
| 5 | 2.747 (2.651, 2.798) | 1.932 (1.889, 1.976) | 1.361 (1.326, 1.384) | 0.962 (0.935, 0.986) | 0.685 (0.665, 0.699) | 0.488 (0.466, 0.494) | 0.347 (0.331, 0.351) | 0.244 (0.235, 0.251) |
| 6 | 1.275 (1.217, 1.99) | 0.972 (0.847, 1.417) | 0.751 (0.608, 0.993) | 0.589 (0.428, 0.711) | 0.454 (0.315, 0.502) | 0.336 (0.24, 0.356) | 0.241 (0.187, 0.253) | 0.17 (0.145, 0.178) |
| 7 | 1.033 (0.993, 1.691) | 0.783 (0.687, 1.184) | 0.596 (0.496, 0.841) | 0.457 (0.353, 0.592) | 0.347 (0.255, 0.422) | 0.264 (0.193, 0.297) | 0.196 (0.149, 0.211) | 0.144 (0.115, 0.15) |
| 8 | 0.933 (0.892, 1.205) | 0.703 (0.623, 0.843) | 0.531 (0.447, 0.603) | 0.39 (0.322, 0.424) | 0.284 (0.232, 0.3) | 0.204 (0.173, 0.21) | 0.146 (0.133, 0.151) | 0.105 (0.098, 0.106) |
| 9 | 0.882 (0.839, 1.028) | 0.64 (0.591, 0.737) | 0.471 (0.421, 0.513) | 0.344 (0.305, 0.372) | 0.241 (0.219, 0.262) | 0.17 (0.155, 0.18) | 0.118 (0.112, 0.13) | 0.082 (0.078, 0.095) |
| 10 | 0.848 (0.813, 0.967) | 0.614 (0.576, 0.708) | 0.45 (0.407, 0.487) | 0.319 (0.289, 0.355) | 0.222 (0.205, 0.248) | 0.157 (0.144, 0.171) | 0.107 (0.102, 0.124) | 0.075 (0.071, 0.091) |
| 11 | 0.845 (0.808, 0.941) | 0.61 (0.574, 0.694) | 0.445 (0.403, 0.475) | 0.314 (0.294, 0.349) | 0.216 (0.211, 0.241) | 0.155 (0.15, 0.167) | 0.107 (0.106, 0.121) | 0.076 (0.074, 0.089) |
| 12 | 0.851 (0.818, 0.941) | 0.613 (0.581, 0.693) | 0.45 (0.407, 0.474) | 0.316 (0.296, 0.349) | 0.224 (0.212, 0.24) | 0.159 (0.154, 0.166) | 0.112 (0.109, 0.121) | 0.078 (0.077, 0.089) |
| 13 | 0.871 (0.841, 0.957) | 0.625 (0.597, 0.703) | 0.462 (0.417, 0.483) | 0.325 (0.303, 0.354) | 0.232 (0.217, 0.244) | 0.164 (0.159, 0.169) | 0.115 (0.113, 0.123) | 0.081 (0.079, 0.091) |
| 14 | 0.903 (0.874, 0.987) | 0.646 (0.62, 0.724) | 0.474 (0.432, 0.498) | 0.334 (0.313, 0.364) | 0.238 (0.224, 0.25) | 0.168 (0.164, 0.174) | 0.118 (0.117, 0.127) | 0.083 (0.081, 0.093) |
| 15 | 0.942 (0.912, 1.026) | 0.673 (0.648, 0.751) | 0.491 (0.452, 0.518) | 0.346 (0.327, 0.377) | 0.241 (0.233, 0.258) | 0.171 (0.169, 0.181) | 0.12 (0.119, 0.132) | 0.085 (0.083, 0.096) |
| 16 | 0.975 (0.955, 1.073) | 0.699 (0.676, 0.784) | 0.509 (0.475, 0.541) | 0.361 (0.34, 0.392) | 0.249 (0.242, 0.269) | 0.179 (0.171, 0.189) | 0.123 (0.12, 0.137) | 0.086 (0.084, 0.1) |
| 17 | 1.025 (0.958, 1.126) | 0.729 (0.678, 0.821) | 0.529 (0.489, 0.568) | 0.369 (0.341, 0.409) | 0.261 (0.242, 0.281) | 0.185 (0.171, 0.198) | 0.125 (0.12, 0.143) | 0.087 (0.084, 0.105) |
| 18 | 1.087 (0.959, 1.184) | 0.769 (0.679, 0.862) | 0.549 (0.491, 0.597) | 0.389 (0.341, 0.427) | 0.271 (0.243, 0.294) | 0.192 (0.172, 0.208) | 0.126 (0.12, 0.15) | 0.087 (0.084, 0.109) |

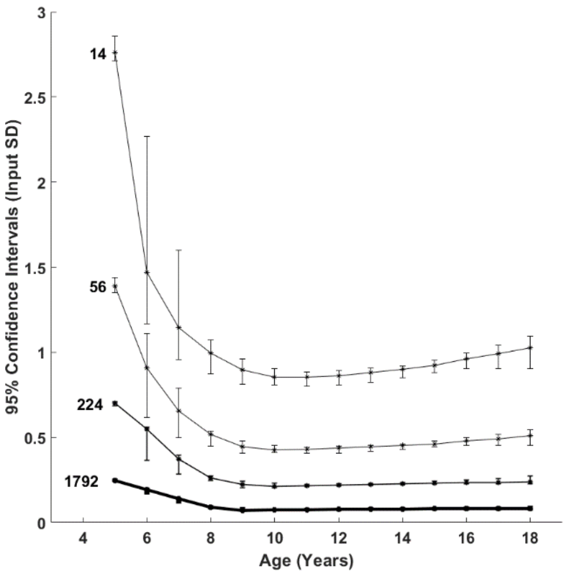


Fig. 5. Span of the 95% CIs across all input SDs for the decay to age 8 input curve. The error bars represent the 2.5 and 97.5 percentiles of the calculated 95% CIs across all input SDs.

#### Decay to age 10

Table 6: CI spans for the decay to age 10 input curve.

| Age | Dataset Size (number of datapoints) | | | | | | | |
|---|---|---|---|---|---|---|---|---|
| | 14 | 28 | 56 | 112 | 224 | 448 | 896 | 1,792 |
| 5 | 2.747 (2.651, 2.798) | 1.932 (1.889, 1.976) | 1.361 (1.326, 1.384) | 0.962 (0.935, 0.986) | 0.685 (0.665, 0.699) | 0.488 (0.466, 0.494) | 0.347 (0.331, 0.351) | 0.244 (0.235, 0.251) |
| 6 | 1.275 (1.217, 1.99) | 0.972 (0.847, 1.417) | 0.751 (0.608, 0.993) | 0.589 (0.428, 0.711) | 0.454 (0.315, 0.502) | 0.336 (0.24, 0.356) | 0.241 (0.187, 0.253) | 0.17 (0.145, 0.178) |
| 7 | 1.033 (0.993, 1.691) | 0.783 (0.687, 1.184) | 0.596 (0.496, 0.841) | 0.457 (0.353, 0.592) | 0.347 (0.255, 0.422) | 0.264 (0.193, 0.297) | 0.196 (0.149, 0.211) | 0.144 (0.115, 0.15) |
| 8 | 0.933 (0.892, 1.205) | 0.703 (0.623, 0.843) | 0.531 (0.447, 0.603) | 0.39 (0.322, 0.424) | 0.284 (0.232, 0.3) | 0.204 (0.173, 0.21) | 0.146 (0.133, 0.151) | 0.105 (0.098, 0.106) |
| 9 | 0.882 (0.839, 1.028) | 0.64 (0.591, 0.737) | 0.471 (0.421, 0.513) | 0.344 (0.305, 0.372) | 0.241 (0.219, 0.262) | 0.17 (0.155, 0.18) | 0.118 (0.112, 0.13) | 0.082 (0.078, 0.095) |
| 10 | 0.848 (0.813, 0.967) | 0.614 (0.576, 0.708) | 0.45 (0.407, 0.487) | 0.319 (0.289, 0.355) | 0.222 (0.205, 0.248) | 0.157 (0.144, 0.171) | 0.107 (0.102, 0.124) | 0.075 (0.071, 0.091) |
| 11 | 0.845 (0.808, 0.941) | 0.61 (0.574, 0.694) | 0.445 (0.403, 0.475) | 0.314 (0.294, 0.349) | 0.216 (0.211, 0.241) | 0.155 (0.15, 0.167) | 0.107 (0.106, 0.121) | 0.076 (0.074, 0.089) |
| 12 | 0.851 (0.818, 0.941) | 0.613 (0.581, 0.693) | 0.45 (0.407, 0.474) | 0.316 (0.296, 0.349) | 0.224 (0.212, 0.24) | 0.159 (0.154, 0.166) | 0.112 (0.109, 0.121) | 0.078 (0.077, 0.089) |
| 13 | 0.871 (0.841, 0.957) | 0.625 (0.597, 0.703) | 0.462 (0.417, 0.483) | 0.325 (0.303, 0.354) | 0.232 (0.217, 0.244) | 0.164 (0.159, 0.169) | 0.115 (0.113, 0.123) | 0.081 (0.079, 0.091) |
| 14 | 0.903 (0.874, 0.987) | 0.646 (0.62, 0.724) | 0.474 (0.432, 0.498) | 0.334 (0.313, 0.364) | 0.238 (0.224, 0.25) | 0.168 (0.164, 0.174) | 0.118 (0.117, 0.127) | 0.083 (0.081, 0.093) |
| 15 | 0.942 (0.912, 1.026) | 0.673 (0.648, 0.751) | 0.491 (0.452, 0.518) | 0.346 (0.327, 0.377) | 0.241 (0.233, 0.258) | 0.171 (0.169, 0.181) | 0.12 (0.119, 0.132) | 0.085 (0.083, 0.096) |
| 16 | 0.975 (0.955, 1.073) | 0.699 (0.676, 0.784) | 0.509 (0.475, 0.541) | 0.361 (0.34, 0.392) | 0.249 (0.242, 0.269) | 0.179 (0.171, 0.189) | 0.123 (0.12, 0.137) | 0.086 (0.084, 0.1) |
| 17 | 1.025 (0.958, 1.126) | 0.729 (0.678, 0.821) | 0.529 (0.489, 0.568) | 0.369 (0.341, 0.409) | 0.261 (0.242, 0.281) | 0.185 (0.171, 0.198) | 0.125 (0.12, 0.143) | 0.087 (0.084, 0.105) |
| 18 | 1.087 (0.959, 1.184) | 0.769 (0.679, 0.862) | 0.549 (0.491, 0.597) | 0.389 (0.341, 0.427) | 0.271 (0.243, 0.294) | 0.192 (0.172, 0.208) | 0.126 (0.12, 0.15) | 0.087 (0.084, 0.109) |

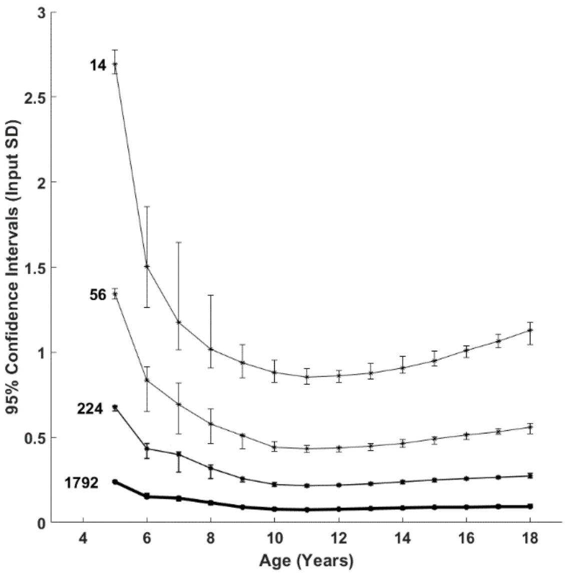


Fig. 6. Span of the 95% CIs across all input SDs for the decay to age 10 input curve. The error bars represent the 2.5 and 97.5 percentiles of the calculated 95% CIs across all input SDs.

#### Decay to age 12

Table 7: CI spans for the decay to age 12 input curve.

| Age | Dataset Size (number of datapoints) | | | | | | | |
|---|---|---|---|---|---|---|---|---|
| | 14 | 28 | 56 | 112 | 224 | 448 | 896 | 1,792 |
| 5 | 2.747 (2.651, 2.798) | 1.932 (1.889, 1.976) | 1.361 (1.326, 1.384) | 0.962 (0.935, 0.986) | 0.685 (0.665, 0.699) | 0.488 (0.466, 0.494) | 0.347 (0.331, 0.351) | 0.244 (0.235, 0.251) |
| 6 | 1.275 (1.217, 1.99) | 0.972 (0.847, 1.417) | 0.751 (0.608, 0.993) | 0.589 (0.428, 0.711) | 0.454 (0.315, 0.502) | 0.336 (0.24, 0.356) | 0.241 (0.187, 0.253) | 0.17 (0.145, 0.178) |
| 7 | 1.033 (0.993, 1.691) | 0.783 (0.687, 1.184) | 0.596 (0.496, 0.841) | 0.457 (0.353, 0.592) | 0.347 (0.255, 0.422) | 0.264 (0.193, 0.297) | 0.196 (0.149, 0.211) | 0.144 (0.115, 0.15) |
| 8 | 0.933 (0.892, 1.205) | 0.703 (0.623, 0.843) | 0.531 (0.447, 0.603) | 0.39 (0.322, 0.424) | 0.284 (0.232, 0.3) | 0.204 (0.173, 0.21) | 0.146 (0.133, 0.151) | 0.105 (0.098, 0.106) |
| 9 | 0.882 (0.839, 1.028) | 0.64 (0.591, 0.737) | 0.471 (0.421, 0.513) | 0.344 (0.305, 0.372) | 0.241 (0.219, 0.262) | 0.17 (0.155, 0.18) | 0.118 (0.112, 0.13) | 0.082 (0.078, 0.095) |
| 10 | 0.848 (0.813, 0.967) | 0.614 (0.576, 0.708) | 0.45 (0.407, 0.487) | 0.319 (0.289, 0.355) | 0.222 (0.205, 0.248) | 0.157 (0.144, 0.171) | 0.107 (0.102, 0.124) | 0.075 (0.071, 0.091) |
| 11 | 0.845 (0.808, 0.941) | 0.61 (0.574, 0.694) | 0.445 (0.403, 0.475) | 0.314 (0.294, 0.349) | 0.216 (0.211, 0.241) | 0.155 (0.15, 0.167) | 0.107 (0.106, 0.121) | 0.076 (0.074, 0.089) |
| 12 | 0.851 (0.818, 0.941) | 0.613 (0.581, 0.693) | 0.45 (0.407, 0.474) | 0.316 (0.296, 0.349) | 0.224 (0.212, 0.24) | 0.159 (0.154, 0.166) | 0.112 (0.109, 0.121) | 0.078 (0.077, 0.089) |
| 13 | 0.871 (0.841, 0.957) | 0.625 (0.597, 0.703) | 0.462 (0.417, 0.483) | 0.325 (0.303, 0.354) | 0.232 (0.217, 0.244) | 0.164 (0.159, 0.169) | 0.115 (0.113, 0.123) | 0.081 (0.079, 0.091) |
| 14 | 0.903 (0.874, 0.987) | 0.646 (0.62, 0.724) | 0.474 (0.432, 0.498) | 0.334 (0.313, 0.364) | 0.238 (0.224, 0.25) | 0.168 (0.164, 0.174) | 0.118 (0.117, 0.127) | 0.083 (0.081, 0.093) |
| 15 | 0.942 (0.912, 1.026) | 0.673 (0.648, 0.751) | 0.491 (0.452, 0.518) | 0.346 (0.327, 0.377) | 0.241 (0.233, 0.258) | 0.171 (0.169, 0.181) | 0.12 (0.119, 0.132) | 0.085 (0.083, 0.096) |
| 16 | 0.975 (0.955, 1.073) | 0.699 (0.676, 0.784) | 0.509 (0.475, 0.541) | 0.361 (0.34, 0.392) | 0.249 (0.242, 0.269) | 0.179 (0.171, 0.189) | 0.123 (0.12, 0.137) | 0.086 (0.084, 0.1) |
| 17 | 1.025 (0.958, 1.126) | 0.729 (0.678, 0.821) | 0.529 (0.489, 0.568) | 0.369 (0.341, 0.409) | 0.261 (0.242, 0.281) | 0.185 (0.171, 0.198) | 0.125 (0.12, 0.143) | 0.087 (0.084, 0.105) |
| 18 | 1.087 (0.959, 1.184) | 0.769 (0.679, 0.862) | 0.549 (0.491, 0.597) | 0.389 (0.341, 0.427) | 0.271 (0.243, 0.294) | 0.192 (0.172, 0.208) | 0.126 (0.12, 0.15) | 0.087 (0.084, 0.109) |

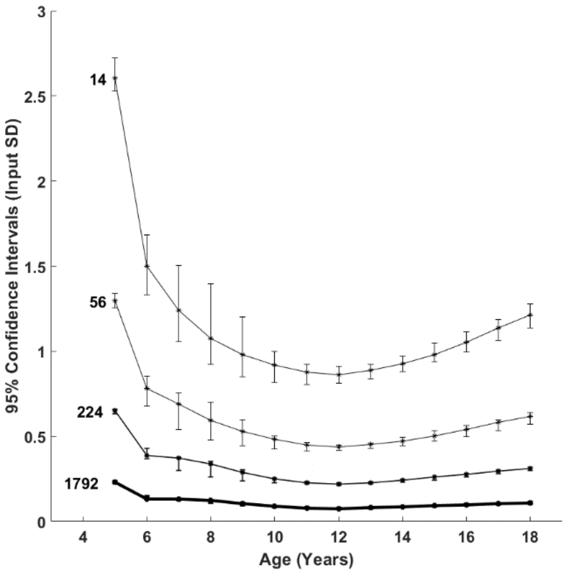


Fig. 7. Span of the 95% CIs across all input SDs for the decay to age 12 input curve. The error bars represent the 2.5 and 97.5 percentiles of the calculated 95% CIs across all input SDs.

#### Decay to age 14

Table 8: CI spans for the decay to age 14 input curve.

| Age | Dataset Size (number of datapoints) | | | | | | | |
|---|---|---|---|---|---|---|---|---|
| | 14 | 28 | 56 | 112 | 224 | 448 | 896 | 1,792 |
| 5 | 2.747 (2.651, 2.798) | 1.932 (1.889, 1.976) | 1.361 (1.326, 1.384) | 0.962 (0.935, 0.986) | 0.685 (0.665, 0.699) | 0.488 (0.466, 0.494) | 0.347 (0.331, 0.351) | 0.244 (0.235, 0.251) |
| 6 | 1.275 (1.217, 1.99) | 0.972 (0.847, 1.417) | 0.751 (0.608, 0.993) | 0.589 (0.428, 0.711) | 0.454 (0.315, 0.502) | 0.336 (0.24, 0.356) | 0.241 (0.187, 0.253) | 0.17 (0.145, 0.178) |
| 7 | 1.033 (0.993, 1.691) | 0.783 (0.687, 1.184) | 0.596 (0.496, 0.841) | 0.457 (0.353, 0.592) | 0.347 (0.255, 0.422) | 0.264 (0.193, 0.297) | 0.196 (0.149, 0.211) | 0.144 (0.115, 0.15) |
| 8 | 0.933 (0.892, 1.205) | 0.703 (0.623, 0.843) | 0.531 (0.447, 0.603) | 0.39 (0.322, 0.424) | 0.284 (0.232, 0.3) | 0.204 (0.173, 0.21) | 0.146 (0.133, 0.151) | 0.105 (0.098, 0.106) |
| 9 | 0.882 (0.839, 1.028) | 0.64 (0.591, 0.737) | 0.471 (0.421, 0.513) | 0.344 (0.305, 0.372) | 0.241 (0.219, 0.262) | 0.17 (0.155, 0.18) | 0.118 (0.112, 0.13) | 0.082 (0.078, 0.095) |
| 10 | 0.848 (0.813, 0.967) | 0.614 (0.576, 0.708) | 0.45 (0.407, 0.487) | 0.319 (0.289, 0.355) | 0.222 (0.205, 0.248) | 0.157 (0.144, 0.171) | 0.107 (0.102, 0.124) | 0.075 (0.071, 0.091) |
| 11 | 0.845 (0.808, 0.941) | 0.61 (0.574, 0.694) | 0.445 (0.403, 0.475) | 0.314 (0.294, 0.349) | 0.216 (0.211, 0.241) | 0.155 (0.15, 0.167) | 0.107 (0.106, 0.121) | 0.076 (0.074, 0.089) |
| 12 | 0.851 (0.818, 0.941) | 0.613 (0.581, 0.693) | 0.45 (0.407, 0.474) | 0.316 (0.296, 0.349) | 0.224 (0.212, 0.24) | 0.159 (0.154, 0.166) | 0.112 (0.109, 0.121) | 0.078 (0.077, 0.089) |
| 13 | 0.871 (0.841, 0.957) | 0.625 (0.597, 0.703) | 0.462 (0.417, 0.483) | 0.325 (0.303, 0.354) | 0.232 (0.217, 0.244) | 0.164 (0.159, 0.169) | 0.115 (0.113, 0.123) | 0.081 (0.079, 0.091) |
| 14 | 0.903 (0.874, 0.987) | 0.646 (0.62, 0.724) | 0.474 (0.432, 0.498) | 0.334 (0.313, 0.364) | 0.238 (0.224, 0.25) | 0.168 (0.164, 0.174) | 0.118 (0.117, 0.127) | 0.083 (0.081, 0.093) |
| 15 | 0.942 (0.912, 1.026) | 0.673 (0.648, 0.751) | 0.491 (0.452, 0.518) | 0.346 (0.327, 0.377) | 0.241 (0.233, 0.258) | 0.171 (0.169, 0.181) | 0.12 (0.119, 0.132) | 0.085 (0.083, 0.096) |
| 16 | 0.975 (0.955, 1.073) | 0.699 (0.676, 0.784) | 0.509 (0.475, 0.541) | 0.361 (0.34, 0.392) | 0.249 (0.242, 0.269) | 0.179 (0.171, 0.189) | 0.123 (0.12, 0.137) | 0.086 (0.084, 0.1) |
| 17 | 1.025 (0.958, 1.126) | 0.729 (0.678, 0.821) | 0.529 (0.489, 0.568) | 0.369 (0.341, 0.409) | 0.261 (0.242, 0.281) | 0.185 (0.171, 0.198) | 0.125 (0.12, 0.143) | 0.087 (0.084, 0.105) |
| 18 | 1.087 (0.959, 1.184) | 0.769 (0.679, 0.862) | 0.549 (0.491, 0.597) | 0.389 (0.341, 0.427) | 0.271 (0.243, 0.294) | 0.192 (0.172, 0.208) | 0.126 (0.12, 0.15) | 0.087 (0.084, 0.109) |

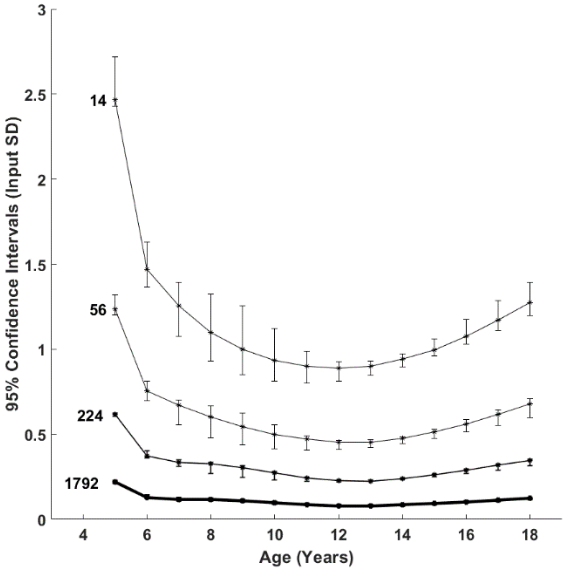


Fig. 8. Span of the 95% CIs across all input SDs for the decaying to age 14 input curve. The error bars represent the 2.5 and 97.5 percentiles of the calculated 95% CIs across all input SDs.

### Non-uniform Datasets

#### Growth Curves

##### Growth to age 8

Table 9: CI spans for the growth to age 8 input curve.

| Age | Dataset size (number of datapoints) | |
|---|---|---|
| | 288 | 576 |
| 5 | 2.84 (2.133, 3.024) | 1.993 (1.517, 2.098) |
| 6 | 0.783 (0.601, 0.904) | 0.582 (0.451, 0.647) |
| 7 | 0.625 (0.426, 0.703) | 0.44 (0.334, 0.524) |
| 8 | 0.49 (0.382, 0.656) | 0.345 (0.292, 0.494) |
| 9 | 0.479 (0.292, 0.641) | 0.336 (0.206, 0.485) |
| 10 | 0.475 (0.203, 0.635) | 0.333 (0.145, 0.482) |
| 11 | 0.474 (0.172, 0.634) | 0.332 (0.122, 0.481) |
| 12 | 0.475 (0.172, 0.634) | 0.332 (0.122, 0.481) |
| 13 | 0.478 (0.177, 0.635) | 0.334 (0.125, 0.482) |
| 14 | 0.482 (0.183, 0.638) | 0.336 (0.129, 0.484) |
| 15 | 0.487 (0.187, 0.642) | 0.339 (0.132, 0.487) |
| 16 | 0.493 (0.189, 0.647) | 0.343 (0.133, 0.489) |
| 17 | 0.5 (0.191, 0.652) | 0.348 (0.134, 0.493) |
| 18 | 0.508 (0.192, 0.658) | 0.352 (0.134, 0.496) |

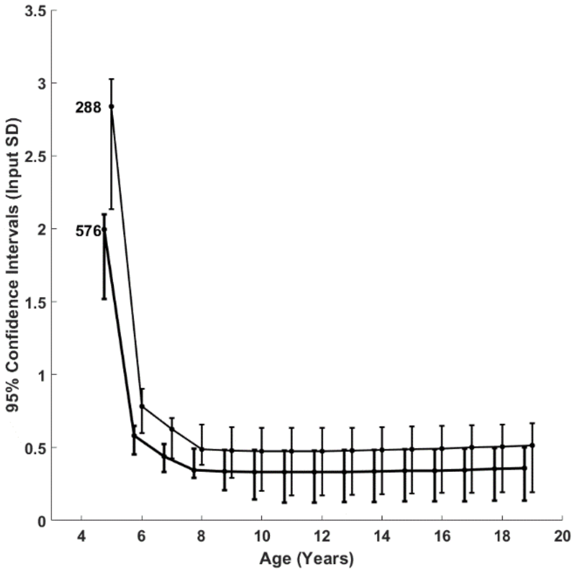


Fig. 9: Span of the 95% CIs across all input SDs for the growth to age 8 input curve. The error bars represent the 2.5 and 97.5 percentiles of the calculated 95% CIs across all input SDs.

#### Growth to age 10

Table 10: CI spans for the growth to age 10 input curve.

| Age | Dataset size (number of datapoints) | |
|---|---|---|
| | 288 | 576 |
| 5 | 2.374 (1.661, 2.623) | 1.541 (1.164, 1.796) |
| 6 | 0.762 (0.64, 0.837) | 0.548 (0.465, 0.586) |
| 7 | 0.569 (0.436, 0.667) | 0.394 (0.332, 0.465) |
| 8 | 0.467 (0.367, 0.621) | 0.323 (0.289, 0.432) |
| 9 | 0.429 (0.338, 0.605) | 0.279 (0.244, 0.421) |
| 10 | 0.421 (0.275, 0.6) | 0.272 (0.193, 0.417) |
| 11 | 0.418 (0.219, 0.6) | 0.269 (0.155, 0.417) |
| 12 | 0.418 (0.182, 0.602) | 0.269 (0.129, 0.418) |
| 13 | 0.421 (0.172, 0.605) | 0.271 (0.123, 0.42) |
| 14 | 0.425 (0.18, 0.61) | 0.275 (0.128, 0.424) |
| 15 | 0.433 (0.194, 0.617) | 0.28 (0.138, 0.428) |
| 16 | 0.442 (0.207, 0.625) | 0.286 (0.148, 0.434) |
| 17 | 0.453 (0.219, 0.634) | 0.294 (0.156, 0.441) |
| 18 | 0.465 (0.227, 0.645) | 0.303 (0.163, 0.449) |

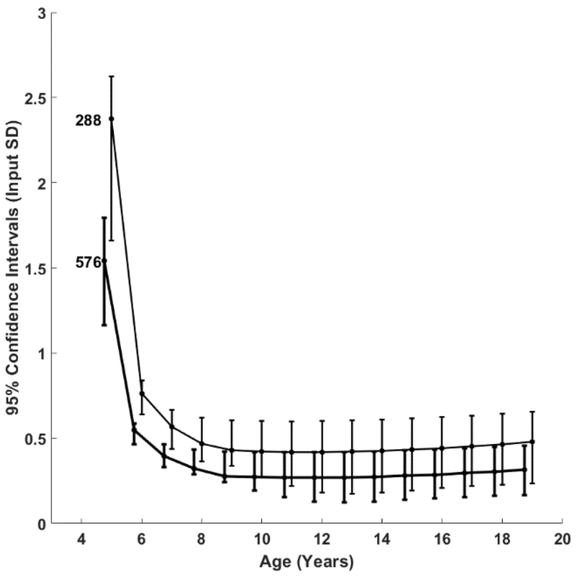


Fig. 10: Span of the 95% CIs across all input SDs for the growth to age 10 input curve. The error bars represent the 2.5 and 97.5 percentiles of the calculated 95% CIs across all input SDs.

#### Growth to age 12

Table 11: CI spans for the growth to age 12 input curve.

| Age | Dataset size (number of datapoints) | |
|---|---|---|
| | 288 | 576 |
| 5 | 1.928 (1.409, 2.401) | 1.254 (0.994, 1.661) |
| 6 | 0.747 (0.655, 0.772) | 0.531 (0.474, 0.539) |
| 7 | 0.5 (0.445, 0.595) | 0.353 (0.327, 0.413) |
| 8 | 0.432 (0.36, 0.542) | 0.297 (0.264, 0.371) |
| 9 | 0.377 (0.319, 0.52) | 0.265 (0.238, 0.352) |
| 10 | 0.354 (0.29, 0.514) | 0.243 (0.208, 0.343) |
| 11 | 0.338 (0.245, 0.514) | 0.238 (0.178, 0.338) |
| 12 | 0.325 (0.21, 0.517) | 0.237 (0.149, 0.339) |
| 13 | 0.319 (0.185, 0.522) | 0.238 (0.129, 0.342) |
| 14 | 0.321 (0.181, 0.53) | 0.243 (0.126, 0.347) |
| 15 | 0.329 (0.196, 0.54) | 0.249 (0.136, 0.353) |
| 16 | 0.344 (0.22, 0.552) | 0.257 (0.154, 0.362) |
| 17 | 0.364 (0.25, 0.566) | 0.268 (0.175, 0.371) |
| 18 | 0.387 (0.276, 0.582) | 0.279 (0.194, 0.383) |

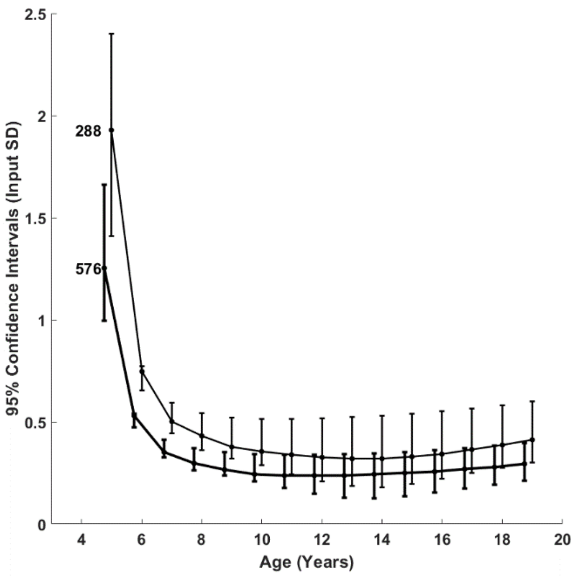


Fig. 11: Span of the 95% CIs across all input SDs for the growth to age 12 input curve. The error bars represent the 2.5 and 97.5 percentiles of the calculated 95% CIs across all input SDs.

#### Growth to age 14

Table 12: CI spans for the growth to age 14 input curve.

| Age | Dataset size (number of datapoints) | |
|---|---|---|
| | 288 | 576 |
| 5 | 1.59 (1.225, 2.182) | 1 (0.861, 1.453) |
| 6 | 0.687 (0.651, 0.748) | 0.492 (0.462, 0.534) |
| 7 | 0.476 (0.452, 0.517) | 0.337 (0.326, 0.368) |
| 8 | 0.384 (0.351, 0.445) | 0.274 (0.249, 0.312) |
| 9 | 0.34 (0.308, 0.408) | 0.241 (0.222, 0.286) |
| 10 | 0.315 (0.274, 0.385) | 0.217 (0.203, 0.273) |
| 11 | 0.302 (0.254, 0.371) | 0.207 (0.179, 0.265) |
| 12 | 0.284 (0.23, 0.364) | 0.2 (0.157, 0.263) |
| 13 | 0.272 (0.202, 0.365) | 0.191 (0.139, 0.264) |
| 14 | 0.271 (0.184, 0.371) | 0.19 (0.13, 0.268) |
| 15 | 0.28 (0.191, 0.382) | 0.197 (0.136, 0.276) |
| 16 | 0.301 (0.222, 0.397) | 0.211 (0.156, 0.286) |
| 17 | 0.331 (0.268, 0.417) | 0.233 (0.187, 0.299) |
| 18 | 0.368 (0.322, 0.441) | 0.254 (0.224, 0.315) |

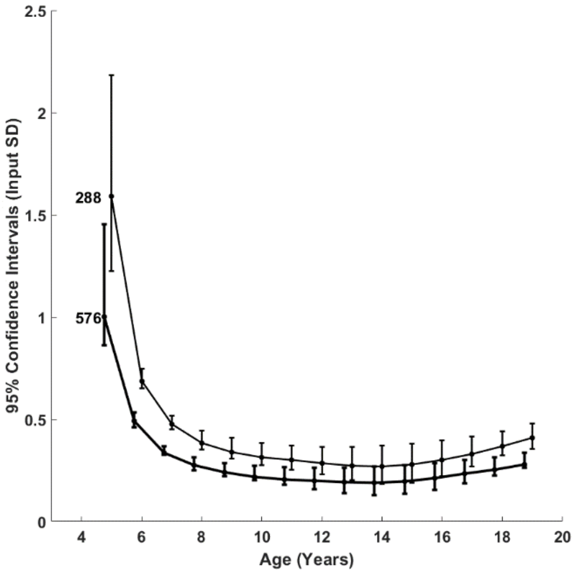


Fig. 12: Span of the 95% CIs across all input SDs for the growth to age 14 input curve. The error bars represent the 2.5 and 97.5 percentiles of the calculated 95% CIs across all input SDs.

#### Decay Curves

##### Decay to age 8

Table 13: CI spans for the decay to age 8 input curve.

| Age | Dataset size (number of datapoints) | |
|---|---|---|
| | 288 | 576 |
| 5 | 3.133 (2.436, 3.243) | 2.122 (1.73, 2.245) |
| 6 | 0.875 (0.646, 0.923) | 0.633 (0.485, 0.664) |
| 7 | 0.578 (0.426, 0.668) | 0.423 (0.324, 0.473) |
| 8 | 0.392 (0.349, 0.506) | 0.278 (0.265, 0.341) |
| 9 | 0.324 (0.225, 0.458) | 0.236 (0.16, 0.309) |
| 10 | 0.302 (0.173, 0.434) | 0.214 (0.123, 0.292) |
| 11 | 0.289 (0.17, 0.422) | 0.202 (0.12, 0.282) |
| 12 | 0.281 (0.174, 0.415) | 0.197 (0.122, 0.277) |
| 13 | 0.278 (0.177, 0.412) | 0.196 (0.124, 0.274) |
| 14 | 0.277 (0.18, 0.411) | 0.197 (0.126, 0.274) |
| 15 | 0.279 (0.18, 0.412) | 0.199 (0.126, 0.274) |
| 16 | 0.283 (0.181, 0.415) | 0.202 (0.126, 0.276) |
| 17 | 0.289 (0.181, 0.418) | 0.205 (0.127, 0.279) |
| 18 | 0.295 (0.181, 0.422) | 0.208 (0.127, 0.283) |

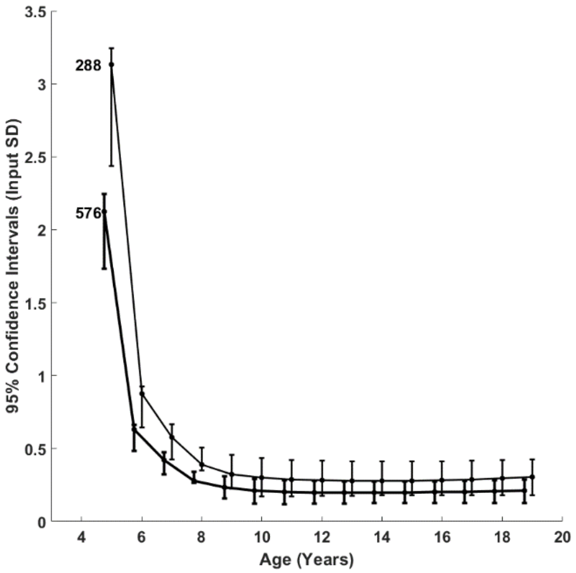


Fig. 13: Span of the 95% CIs across all input SDs for the decay to age 8 input curve. The error bars represent the 2.5 and 97.5 percentiles of the calculated 95% CIs across all input SDs.

#### Decay to age 10

Table 14: CI spans for the decay to age 10 input curve.

| Age | Dataset size (number of datapoints) | |
|---|---|---|
| | 288 | 576 |
| 5 | 2.422 (1.879, 2.869) | 1.534 (1.323, 1.975) |
| 6 | 0.81 (0.698, 0.859) | 0.564 (0.514, 0.608) |
| 7 | 0.576 (0.472, 0.596) | 0.41 (0.362, 0.424) |
| 8 | 0.447 (0.38, 0.473) | 0.321 (0.292, 0.338) |
| 9 | 0.344 (0.327, 0.414) | 0.243 (0.231, 0.291) |
| 10 | 0.297 (0.245, 0.381) | 0.183 (0.173, 0.267) |
| 11 | 0.267 (0.189, 0.361) | 0.155 (0.134, 0.252) |
| 12 | 0.251 (0.172, 0.348) | 0.141 (0.122, 0.243) |
| 13 | 0.245 (0.176, 0.34) | 0.138 (0.125, 0.238) |
| 14 | 0.245 (0.184, 0.338) | 0.14 (0.13, 0.236) |
| 15 | 0.25 (0.192, 0.339) | 0.146 (0.135, 0.237) |
| 16 | 0.258 (0.199, 0.344) | 0.153 (0.141, 0.24) |
| 17 | 0.268 (0.203, 0.351) | 0.16 (0.144, 0.245) |
| 18 | 0.28 (0.205, 0.361) | 0.167 (0.146, 0.252) |

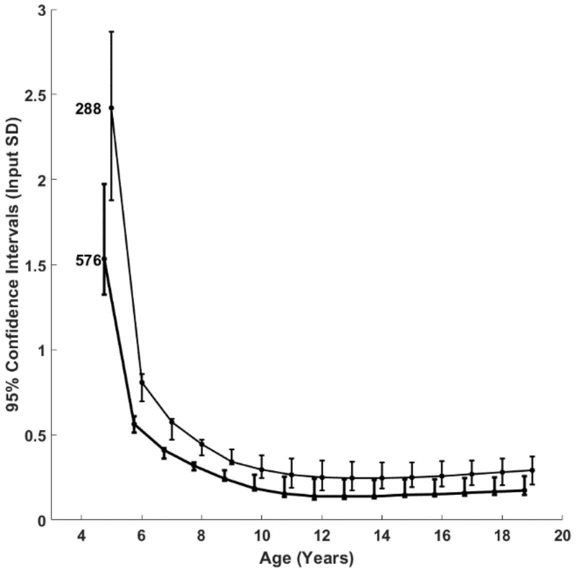


Fig. 14: Span of the 95% CIs across all input SDs for the decay to age 10 input curve. The error bars represent the 2.5 and 97.5 percentiles of the calculated 95% CIs across all input SDs.

#### Decay to age 12

Table 15: CI spans for the decay to age 12 input curve.

| Age | Dataset size (number of datapoints) | |
|---|---|---|
| | 288 | 576 |
| 5 | 1.939 (1.579, 2.556) | 1.208 (1.125, 1.708) |
| 6 | 0.772 (0.716, 0.852) | 0.544 (0.526, 0.603) |
| 7 | 0.525 (0.48, 0.616) | 0.368 (0.338, 0.441) |
| 8 | 0.423 (0.394, 0.505) | 0.298 (0.292, 0.363) |
| 9 | 0.359 (0.341, 0.443) | 0.254 (0.246, 0.317) |
| 10 | 0.297 (0.282, 0.403) | 0.207 (0.2, 0.288) |
| 11 | 0.256 (0.23, 0.377) | 0.166 (0.161, 0.269) |
| 12 | 0.229 (0.189, 0.359) | 0.14 (0.135, 0.256) |
| 13 | 0.216 (0.174, 0.348) | 0.129 (0.124, 0.249) |
| 14 | 0.214 (0.18, 0.344) | 0.131 (0.127, 0.245) |
| 15 | 0.222 (0.195, 0.345) | 0.141 (0.136, 0.245) |
| 16 | 0.236 (0.211, 0.35) | 0.153 (0.147, 0.249) |
| 17 | 0.256 (0.226, 0.36) | 0.166 (0.158, 0.255) |
| 18 | 0.278 (0.238, 0.373) | 0.179 (0.167, 0.264) |

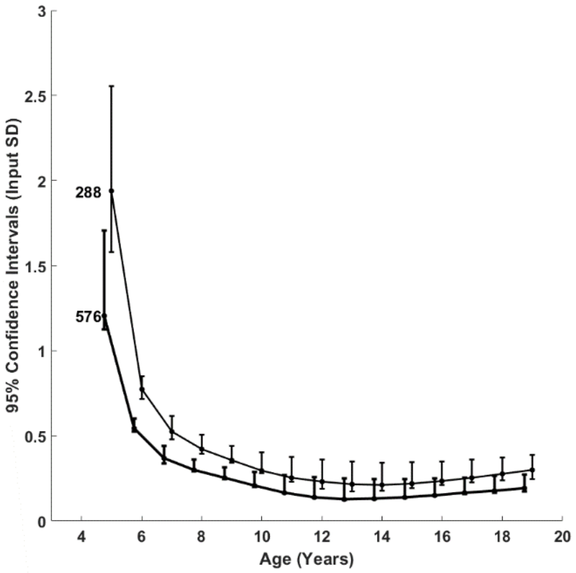


Fig. 15: Span of the 95% CIs across all input SDs for the decay to age 12 input curve. The error bars represent the 2.5 and 97.5 percentiles of the calculated 95% CIs across all input SDs.

#### Decay to age 14

Table 16: CI spans for the decay to age 14 input curve.

| Age | Dataset size (number of datapoints) | |
|---|---|---|
| | 288 | 576 |
| 5 | 1.53 (1.386, 2.329) | 1.019 (0.979, 1.513) |
| 6 | 0.752 (0.727, 0.8) | 0.534 (0.525, 0.559) |
| 7 | 0.497 (0.464, 0.591) | 0.337 (0.328, 0.413) |
| 8 | 0.395 (0.373, 0.486) | 0.273 (0.262, 0.343) |
| 9 | 0.344 (0.336, 0.421) | 0.241 (0.237, 0.3) |
| 10 | 0.306 (0.288, 0.376) | 0.213 (0.205, 0.27) |
| 11 | 0.261 (0.246, 0.344) | 0.182 (0.175, 0.249) |
| 12 | 0.219 (0.212, 0.321) | 0.153 (0.15, 0.233) |
| 13 | 0.193 (0.187, 0.307) | 0.134 (0.131, 0.224) |
| 14 | 0.184 (0.18, 0.301) | 0.129 (0.127, 0.219) |
| 15 | 0.196 (0.193, 0.301) | 0.138 (0.136, 0.218) |
| 16 | 0.223 (0.217, 0.309) | 0.156 (0.152, 0.223) |
| 17 | 0.254 (0.249, 0.323) | 0.179 (0.174, 0.231) |
| 18 | 0.289 (0.277, 0.342) | 0.203 (0.196, 0.243) |

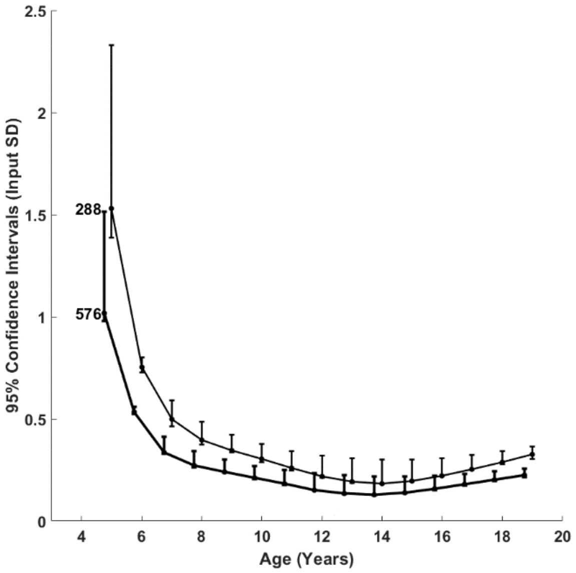


Fig. 16: Span of the 95% CIs across all input SDs for the decay to age 14 input curve. The error bars represent the 2.5 and 97.5 percentiles of the calculated 95% CIs across all input SDs.
